# Elucidating the activation mechanism of the proton-sensing GPR68 receptor

**DOI:** 10.1101/2023.12.08.570878

**Authors:** Christos Matsingos, Lesley A. Howell, Peter J. McCormick, Arianna Fornili

## Abstract

GPR68 is a proton-sensing G-protein Coupled Receptor (GPCR) involved in a variety of physiological processes and disorders including neoplastic pathologies. While GPR68 and few other GPCRs have been shown to be activated by a decrease in the extracellular pH, the molecular mechanism of their activation remains largely unknown. In this work, we used a combined computational and *in vitro* approach to provide new insight into the activation mechanism of the receptor. Molecular Dynamics simulations of GPR68 were used to model the changes in residue interactions and motions triggered by pH. Global and local rearrangements consistent with partial activation were observed upon protonation of the inactive state. Selected extracellular histidine and transmembrane acidic residues were found to have significantly upshifted p*K*_a_ values during the simulations, consistently with their previously hypothesised role in activation through changes in protonation state. Moreover, a novel pairing between histidine and acidic residues in the extracellular region was highlighted by both sequence analyses and simulation data and tested through site-directed mutagenesis. At last, we identified a previously unknown hydrophobic lock in the extracellular region that might stabilise the inactive conformation and regulate the transition to the active state.g

## INTRODUCTION

GPR68 is part of a small group of G protein-coupled receptors (GPCRs) that are activated by a decrease in pH (‘proton-sensing’)^1,2^. This receptor is coupled with different types of G-proteins^1,3,4^ and is involved in a variety of biological processes including cellular differentiation in bone tissue^5,6^, regulation of insulin levels^7^, mechanosensing^8,9^, hippocampal neurogenesis^10^, memory^11^ and hormone release from the pituitary gland^12,13^. GPR68 has also been shown to have a role in tumour growth^4,14–18^ and metastasis^19,20^, and has been found in many neoplastic tissues^21^, possibly associated with the microacidic environment involved in solid tumours^22^.

No experimental structure of GPR68 or closely related protein has been published so far and the molecular details of the proton-sensing mechanism in GPCRs are still unclear. After the discovery of GPR4 and GPR68 as proton-sensing GPCRs in 2003^1^, homology models and mutagenesis studies initially indicated that a cluster of histidine residues located on the extracellular side of these receptors (green in Figure 1) was mostly responsible for pH sensitivity^1,11,23^. Upon a decrease in environmental pH, histidines in the cluster would become protonated, initiating activation through a series of unknown conformational changes. Mutagenesis showed that these histidines have different degrees of involvement in proton-sensing^23^, ranging from H20^1.31^ and H169^EL2^, which have the most consistent effects on pH sensitivity, to H89^EL1^, H159^4.63^ and H175^EL2^, which show a significant effect only when mutated in combination with other histidines. The two transmembrane H245^6.52^ and H269^7.36^ residues (purple in Figure 1), while significantly affecting proton sensitivity when mutated to phenylalanine, were found to lead to modest effects when mutated to alanine, suggesting they have a structural role rather than a direct involvement in proton-sensing through changes in their protonation state.

**Figure 1.**
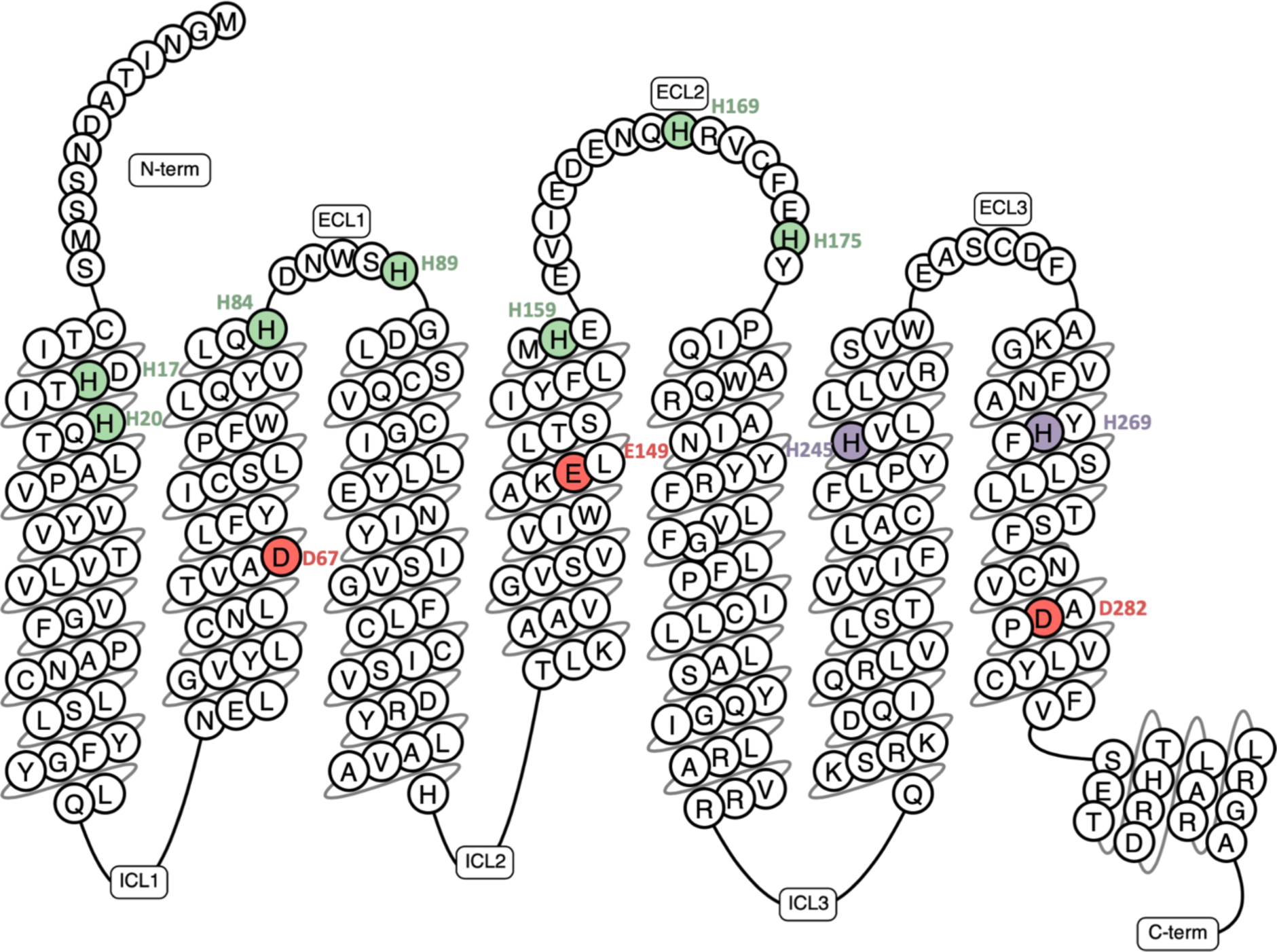
Snake plot of GPR68 from GPCRdb^25^. Extracellular (green) and transmembrane (purple) histidine residues are highlighted with colour, together with the acidic triad (red).

Subsequent studies placed emphasis on the role of acidic residues in proton sensitivity. Evolutionary analyses identified an ‘acidic triad’ composed of residues D67^2.50^, E149^4.53^ and D282^7.49^ (red in Figure 1), which is conserved in the proton-sensing GPR4, GPR65 and GPR68 receptors and was shown to affect their activity upon mutation^2^. Moreover, a recent computational study predicted the existence of a pH-dependent network of water-mediated hydrogen bond interactions connecting histidine and acidic residues in the extracellular and transmembrane regions of GPR68^24^.

In this work, we provide new insight into the activation mechanism of GPR68 through Molecular Dynamics simulations of modelled inactive and active states of the receptor and through mutagenesis studies. A sequence of protonation events is shown to lead to the beginning of the transition towards the active state. Our findings not only confirm the importance of the acidic triad in the activation mechanism but also lead to the identification of key pairings between acidic and histidine residues and of a previously unknown hydrophobic lock in the extracellular region.

## RESULTS AND DISCUSSION

### Protonation of key residues affects inter-residue distances

MD simulations were used to model the structural changes triggered by protonation of the inactive state of GPR68. Simulations were first carried out in three replicas starting from an inactive state model (see Methods section) with all the residues in their standard protonation state (‘U’ trajectories in Table S1). The predicted p*K*_a_ values of histidine and acidic amino acids were calculated for each trajectory frame in order to estimate the propensity of these residues to become protonated within the activation window of GPR68. Interestingly, the p*K*_a_ values of H17^1.28^, H84^2.67^ and H169^EL2^ were found to be ≥ 6.8 (proton EC_50_ in experiments measuring cAMP accumulation in transiently transfected HEK293T cells^23^) for at least part of the trajectories (Figures 2A and S1). These residues are part of the group of histidines considered to be involved in proton-sensing (Table S2). Our data show that among the extracellular histidines, H169^EL2^ seems to be the most likely to get protonated, at least in the initial stages of the activation. The two transmembrane residues H245^6.52^ and H269^7.36^ have p*K*_a_ values consistently low (< 6) for all (H245^6.52^) or most (H269^7.36^) of the simulated time, confirming the indication from previous experiments^23^ that these histidines have more a structural role than a proton-sensing one. Among the acidic residues, the DyaD residues D67^2.50^ and D282^7.49^ and the apEx residue E149^4.53^ were all found to have p*K*_a_ values shifted upwards from their standard reference values (3.8 for aspartates and 4.8 for glutamates), with D67^2.50^ having a p*K*_a_ ≥ 6.8 for at least 60% of the trajectory in all the replicas (Figure 2A).

**Figure 2.**
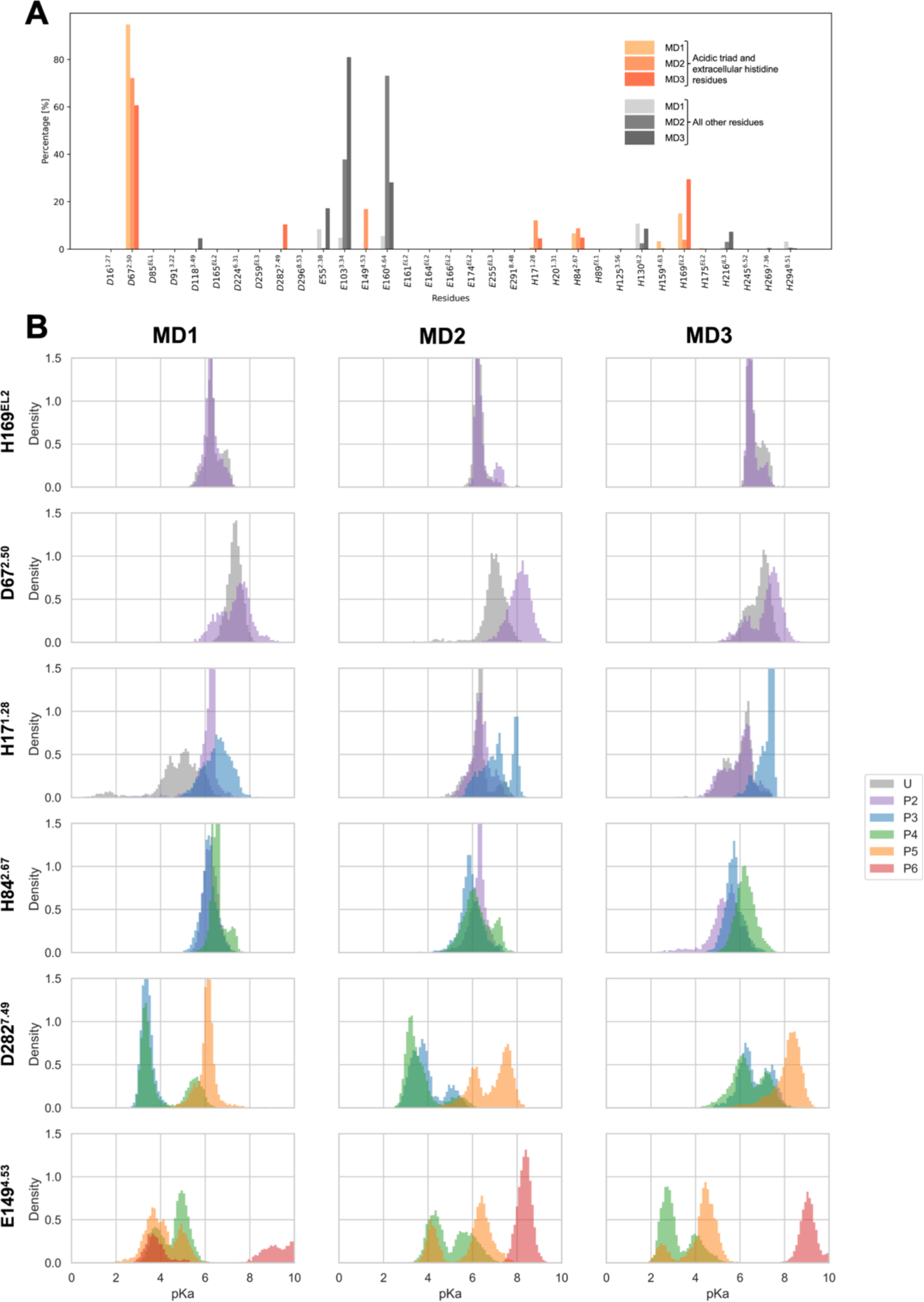
Predicted p*K*_a_ values of histidine and acidic residues during the MD simulations of GPR68. A) Percentage of frames with p*K*_a_ values ≥ 6.8 for each of the acidic and histidine residues in GPR68. Values were calculated from the 300-ns production runs of the ‘U’ trajectories. The residues forming the acidic triad and the extracellular histidine residues are shown with orange shades. B) Distribution of p*K*_a_ values for the residues selected to change their protonation state during the MD simulations. For each residue, p*K*_a_ distributions are shown for the production run of the trajectory where the residue was protonated, together with one (for D67^2.50^ and H169^EL2^) or two (for all the other residues) trajectories before in the ‘U’ to ‘P6’ protonation sequence.

A sequence of protonation events (Table S1) was designed based on the distribution of p*K*_a_ values of selected residues. For each ‘U’ replica, a protonation event was the starting point of a new simulation, which was used to recalculate the distribution of p*K*_a_ values. Residues were selected to be protonated based on whether they have been previously found through experiments to be involved in proton-sensing (acidic triad + proton-sensing histidines in Table S2) and whether they showed a consistently high p*K*_a_ value distribution in our simulations. Consistently with the analysis of the ‘U’ set of replicas described above, D67^2.50^ and H169^EL2^ were protonated at first (‘P2’ trajectories). While the p*K*_a_ ranges of H17^1.28^ and H84^2.67^ were found to be comparable in ‘P2’, H17^1.28^ showed narrower distributions at higher values compared to H84^2.67^ (purple in Figure 2B). Moreover, the p*K*_a_ of H17^1.28^ seemed to be more sensitive to D67^2.50^ and H169^EL2^ protonation as its values were consistently high in all replicas only after these residues were protonated, so H17^1.28^ was protonated next (‘P3’ replicas), followed by H84^2.67^ (‘P4’). In the last two stages, the two remaining residues of the acidic triad were protonated. The DyaD residue D282^7.49^ showed high p*K*_a_ values (> 6.8) in at least one of the ‘P4’ replicas (green in Figure 2B) and it was simulated as protonated in the ‘P5’ trajectories. This led to an upward shift of the p*K*_a_ of the conserved apEx residue E149^4.53^ (orange in Figure 2B), which was then protonated in the last set of replicas (‘P6’).

The different protonation states were simulated for varying times (Table S1) to allow for the equilibration of p*K*_a_ values and for the Root Mean Square Deviation (RMSD) of the Cα atoms to reach a steady state. For each set of simulations, the ‘U’ and ‘P2’ to ‘P6’ trajectories were concatenated together, to give three pseudo-trajectories (MD1 to MD3). It is interesting to note that protonating a residue often promoted a further increase of its p*K*_a_, which is particularly evident for H17^1.28^, D282^7.49^ and E149^4.53^ (Figure 2B). This is most likely due to the protonated form of the residues moving into a more favourable environment, e.g. closer to amino acids with opposite charge or that can form new favourable interactions.

The pseudo-trajectories were analysed to identify possible changes in inter-residue interactions induced by protonation, such as the disruption of the previously hypothesised histidine cluster on the extracellular side of the receptor^1,23^. Indeed, monitoring the distance between pairs of extracellular histidines showed that as the number of protonated residues increases, the H17^1.28^-H169^EL2^ and H84^2.67^-H169^EL2^ distances increase in two out of our three pseudo-trajectories (MD1 and MD2 in Figure S2). However, distance values below 5 Å were rarely found for those pairs in all the trajectories, even before protonation. It is thus unlikely that the movement of these histidine residues away from each other is driven by electrostatic repulsion, but rather, a more complex process is taking place.

A closer look at the environment of H17^1.28^, H84^2.67^ and H169^EL2^ shows that they are in close spatial and sequence proximity to acidic residues. Residue H17^1.28^ is preceded in the sequence by D16^1.27^, H84^2.67^ by D85^EL1^ and H169^EL2^ by the triplet of residues E164^EL2^, D165^EL2^ and E166^EL2^, which will be referred to in the following as the EDE segment. These acidic residues are expected to promote high p*K*_a_ values for the histidine residues especially when they form direct interactions with them. The D16^1.27^/H17^1.28^ and H84^2.67^/D85^EL1^ sequence pairs were found to form alternating inter- and intra-pair interactions during the MD2 pseudo-trajectory depending on the histidine protonation state (Figure 3). In the ‘P3’ part, D85^EL1^ starts forming inter-pair interactions with the newly protonated H17^1.28^ (I and magenta line in Figure 3), to then switch to intra-pair interactions with H84^2.67^ (II and brown line) once the latter is protonated (‘P4’). At the same time, H17^1.28^ switches to intra-pair interactions with D16^1.27^ (II and yellow line). This alternating pattern of inter- and intra-pair interactions was repeatedly found during MD2 (e.g. IV and V). Intra-pair interactions were also observed between H169^EL2^ and the EDE residues towards the end of the pseudo-trajectory (III and VI), after D282^7.49^ protonation (‘P5’). While the specific sequence of interactions was different in the other two pseudo-trajectories, all the discussed intra-pair interactions were found in MD1 and MD3, and the inter-pair interaction H17^1.28^-D85^EL1^ was observed again in MD3 (Figures S3 and S4). Interestingly, stable interactions between histidines and acidic residues were found only upon histidine protonation, suggesting that they might be part of the conformational changes that take place during activation.

**Figure 3.**
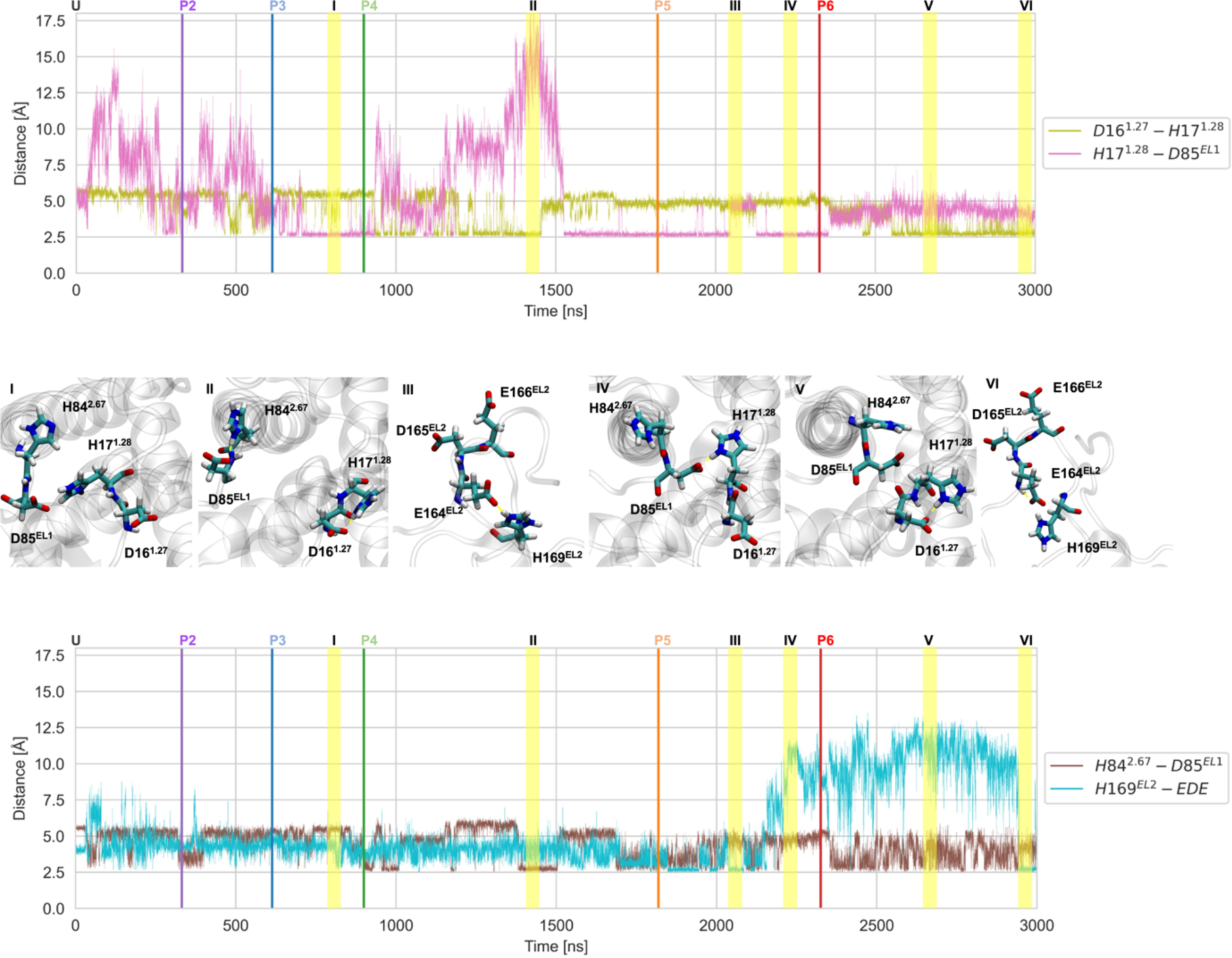
Time evolution of selected distances between histidine and acidic residues during the MD2 pseudo-trajectory. The minimum distance calculated over all possible pairs of side chain heavy atoms is plotted for the D16^1.27^-H17^1.28^ and H17^1.28^-D85^EL1^ (top panel) and H84^2.67^- D85^EL1^ and H169^EL2^-EDE (bottom panel) pairs. Vertical lines separate the individual trajectories with different protonation states (‘U’ to ‘P6’). A close-up view of the residue pairs (sticks) is shown in the middle panel for representative time points I-VI (marked in yellow in the distance plots). Dashed yellow lines indicate hydrogen bond interactions.

### Pairing of histidine and acidic residues is a shared trait among proton-sensing GPCRs

A sequence analysis was performed in order to assess the significance and frequency of the histidine-acidic residue pairing in GPR68 and other proton-sensing GPCRs (GPR4, GPR65 and GPR132). For each receptor, all histidine residues were first classified into five categories according to their position in the sequence (‘extracellular’, ‘intracellular’ or ‘transmembrane’) and, for extracellular histidines, whether they have been experimentally tested and found to be involved in proton-sensing (Table S2 and Figure 4A). Homologous sequences of each receptor were then collected and aligned (see Methods). Analysis of the multiple sequence alignments showed a higher propensity for extracellular histidine residues to have a glutamate or aspartate residue within two positions in the sequence compared not only to transmembrane histidines but also intracellular ones (Figure 4A). On average, this propensity seems to be higher for extracellular histidines considered to be involved in proton-sensing (green) compared to those not involved (blue).

**Figure 4.**
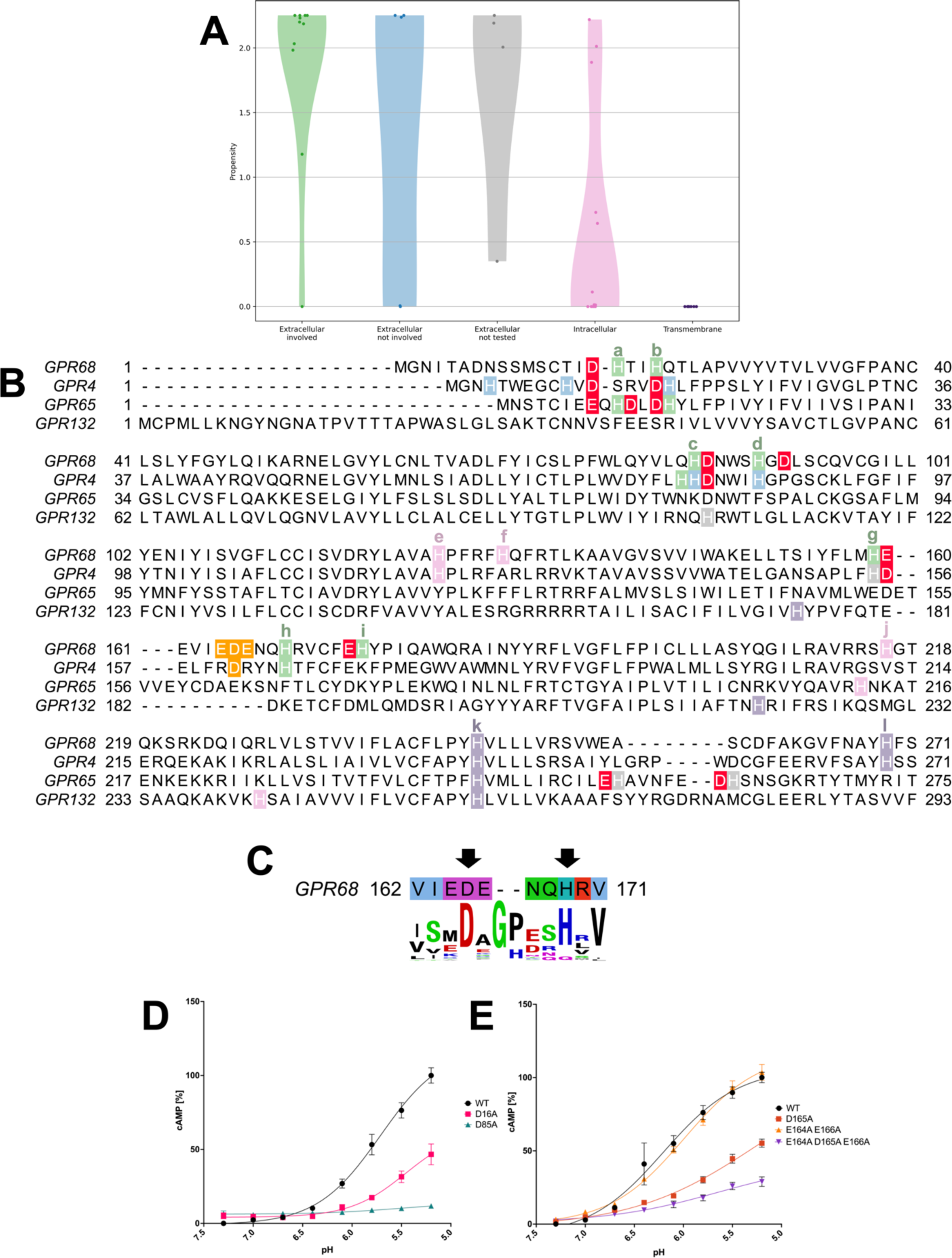
Importance of acidic residues in proton-sensing GPCRs. A) Violin plots showing the propensity of histidine residues in proton-sensing GPCRs to have an acidic residue within two positions in the sequence as calculated from multiple sequence alignments for each receptor.

These findings are exemplified by the human sequences of the receptors (Figure 4B). For GPR68, in addition to the H17^1.28^ (a in the figure), H84^2.67^ (c) and H169^EL2^ (h) residues discussed in the previous section, also the extracellular H89^EL1^ (d), H159^4.63^ (g) and H175^EL2^ (i) histidines, which are thought to have a role in proton-sensing even if secondary (Table S2), are found in close proximity of an acidic residue (D91^3.22^, E160^4.64^ and E174^EL2^ respectively). While H169^EL2^ has a relatively high propensity value (1.98), in the human sequence it is three positions apart from the first acidic residue (E166^EL2^). On the other hand, H169^EL2^ is close in space to two additional acidic residues from the EDE segment, one of which (D165^EL2^) is very well conserved within GPR68 homologs (Figure 4C). Residues H245^6.52^ (k) and H269^7.36^ (l) in Figure 4B do not have any acidic amino acids in their vicinity, consistently with their location in the transmembrane region and their lower probability of being involved in proton-sensing through changes of their protonation state^23^.

The pairing of acidic residues with extracellular histidines extends to other proton-sensing GPCRs. In GPR65, both proton-sensing^26^ residues H10^NTER^ (a) and H14^1.32^ (1 position after b) show a high propensity of having acidic residues nearby (>2.18), with H10^NTER^ (H17^1.28^ in GPR68) being sandwiched between two acidic residues in the human sequence. A high propensity (>2) was also found for the two extracellular histidines H254^6.63^ and H261^7.24^, which however have not been experimentally tested so far for proton-sensing. In human GPR4, both extracellular proton-sensing^27^ residues H79^2.66^ (close to H84^2.67^ in GPR68) and H165^EL2^ (H169^EL2^ in GPR68) are close to an acidic residue in the sequence, with D161^EL2^ possibly taking a similar role as D165^EL2^ in GPR68. GPR132 lacks extracellular histidine residues in the same positions as the other receptors except for H106^EL1^ (1 position after c), for which no significant pairing with acidic residues was found (propensity = 0.35). Evidence of proton-sensing of this receptor remains limited^2^.

Both our MD simulations and sequence analysis highlight a possible role for D16^1.27^, D85^EL1^ and the EDE segment in the activation of GPR68. These findings were tested through site-directed mutagenesis, where these residues were mutated to alanine (individually or in combination) and the production of cAMP was measured using a real-time biosensor. Both the D16A and D85A mutants show a significant reduction in activity compared to the wild type (Figure 4D), which confirms their involvement in the activation mechanism. An almost complete loss of activity was found for D85A while some was preserved in D16A. This is consistent with the larger range of interactions observed for D85^EL1^ during the simulations compared to D16^1.27^, as the residue switches interaction partners according to their protonation state and seems to have a more central role in the rearrangement of proton-sensing histidine residues. In the EDE segment, a large drop in activity was observed for D165A (red in Figure 4E). While mutating both E164^EL2^ and E166^EL2^ (orange) did not affect activity, a significant drop was observed for the E164A/D165A/E166A triple mutant compared to D165A (purple), suggesting that the two glutamate residues are not necessary for the function of the receptor in the wild type sequence but can at least partially assume the role of D165^EL2^ in the activation process when the latter is mutated to alanine.

In summary, we found that the pairing of acidic and histidine residues is a recurring feature in proton-sensing GPCRs, which is more likely to occur on the extracellular side of the receptor and when histidines are important for proton-sensing. Our simulations, sequence analysis and mutagenesis data support the involvement of D16^1.27^, D85^EL1^ and the EDE segment in the activation mechanism of GPR68. The presence of these acidic residues close to proton-sensing histidines could contribute to the modulation both of their p*K*_a_ and their arrangement as shown in the previous section. This is not unique to proton-sensing GPCRs as similar roles are undertaken for example by aspartate residues in the catalytic triad of serine proteases^28^.

The results for all histidine residues of GPR68, GPR4, GPR65 and GPR132 are shown. Histidine residues are classified as transmembrane, extracellular or intracellular, with the extracellular ones further divided into involved in proton-sensing, not involved or not tested (Table S2). B) Multiple sequence alignment (GPR68 residues 1-271) of proton-sensing GPCRs as generated by PRALINE^29^. Histidine residues are colour-coded as in panel A. Acidic residues within 2 positions of histidines are highlighted in red, while the EDE segment in GPR68 and acidic residue at equivalent positions in GPR4 (D161^EL2^) are highlighted in orange. Lowercase letters are used to label GPR68 histidine residues. C) Sequence logo of the 163-171 region in GPR68 generated with WebLogo^30^ using the multiple sequence alignment described in the main text. D/E) pH concentration-response curves for cAMP production in HEK293 cells for the mutants D16A and D85A (D panel) and D165A and E164A/E166A and E154A/D165A/E166A (E panel) compared to wild type. Data represent the mean ± SEM from *n* = 3 individual replicates performed in triplicate. The error bars that are not visible lie within the dimensions of the symbol.

### Protonation leads to movement of TM6 and TM7 in GPR68

The simulations performed at increasing degrees of protonation (‘U’ to ‘P6’) were further analysed to detect possible evidence of transition towards the active state. A collective variable to monitor the transition was first identified by performing a principal component analysis (PCA) on the pseudo-trajectory composed of the ‘U’ inactive state and active state replicas (Cα atoms of the transmembrane helices only). The first principal component (PC1) was found to account for almost half of the total variance (48.2%, Table S3) and individually projecting the replicas over it showed a clear separation between the inactive and active state (Figure S5), indicating that PC1 is suitable to monitor the progression of simulations along the inactive to active transition. The corresponding pseudo-motion involves mostly the outward movement of TM6 and an inward movement of TM7 (Figure 5A and B), consistently with what is expected for class A GPCR activation^31^. Remarkably, the projection of the ‘U’ to ‘P6’ simulations over PC1 showed that in two out of the three sets of replicas (MD1 and MD2 pseudo-trajectories) the system gets closer to the active state as the number of protonated residues increases (Figure 5C and S6).

**Figure 5.**
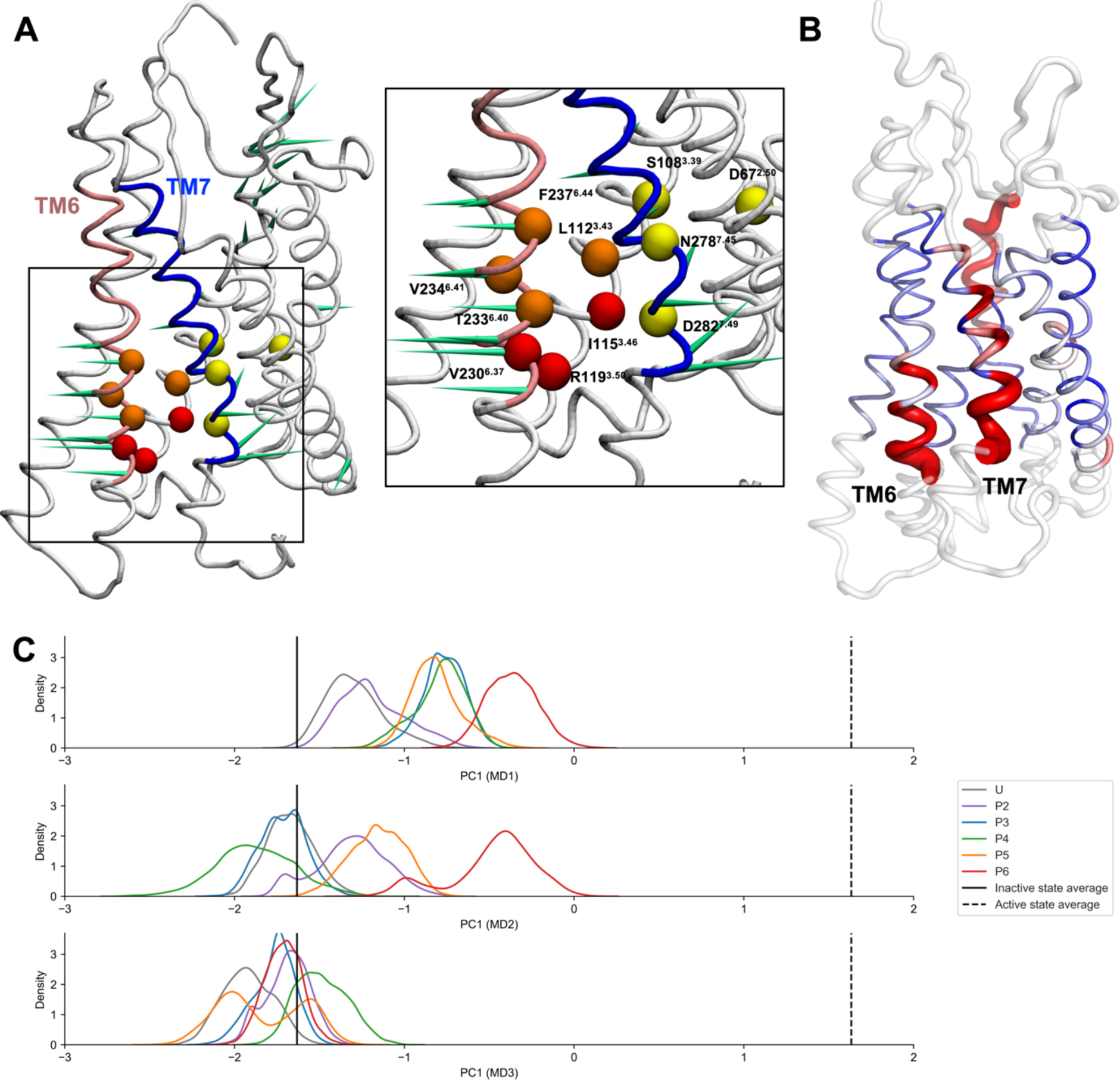
Principal Component Analysis of GPR68 simulations. A) Porcupine representation of PC1. The cyan spikes show the direction and relative amplitude of motion of each residue along PC1. Only spikes with length ≥ 30% of the maximum value are shown. Colour is used to highlight TM6 (pink) and TM7 (blue), while the Cα atoms of the residues involved in the allosteric Na^+^ binding site (yellow), in the hydrophobic lock (orange) and the microswitch residues of TM6 (red) are shown as spheres. Residues are labelled in the close-up view. B) Structural mapping of the Cα RMSF profile for the motion along PC1 observed in the MD2 pseudo-trajectory. RMSF values are colour-mapped from blue to red onto a representative structure of the inactive state (centroid of the most populated cluster of ‘U’ replicas). The thickness of the tube representation is proportional to the RMSF value. Extra- and intracellular loops are shown as cartoon (clear) without RMSF values assigned to them. C) Histograms of the projection of the MD1, MD2 and MD3 pseudo-trajectories (‘U’ to ‘P6’ simulations) on PC1 (the initial equilibration of the ‘U’ trajectories is not included). Vertical lines indicate the mean values of PC1 calculated over the ‘U’ inactive (solid black line) and active (dashed black line) state trajectories.

The pseudomotion described by PC1 is more prominent in the parts of TM6 and TM7 helices closer to the intracellular side (Figure 5B). These regions include three groups of microswitch residues (Figure 6.), whose spatial rearrangements are usually indicative of class A GPCR activation^31^. The hydrophobic lock microswitch includes residues at positions 3.43, 6.41, 6.40 and 6.44^32^ (orange spheres in Figure 5A and orange shading in Figure 6.). In class A GPCRs, the outward movement of the cytoplasmic end of TM6 is associated with an opening of the hydrophobic lock^31,32^. Residues at positions 6.40, 6.44, and, to a lesser extent, 6.41 on TM6 form a hydrophobic cage around L3.43 in the inactive state^32^. During activation, contacts with this residue decrease allowing the movement of TM6. The distances between L112^3.43^ and each of the other lock residues were monitored in our ‘U’ to ‘P6’ simulations to identify possible changes in the hydrophobic lock upon protonation. An increase of the L112^3.43^-F237^6.44^ and L112^3.43^-V234^6.41^ distances was observed in MD1 and MD2 (Figure S7), indicating an opening of the lock (orange-lined close-up view in Figure 6.). MD3 showed a decrease or no significant changes in these distances, congruent with no activation taking place in this set of replicas. No consistent results were found for the L112^3.43^-T233^6.40^ distance across the replicas. The discrepancy from the expected behaviour could be due to the fact that the residue at position 6.40 is often a hydrophobic residue in other class A GPCRs^32^. A threonine in this position would not form extensive hydrophobic interactions so that local rearrangements of this residue could occur independently from activation.

**Figure 6.**
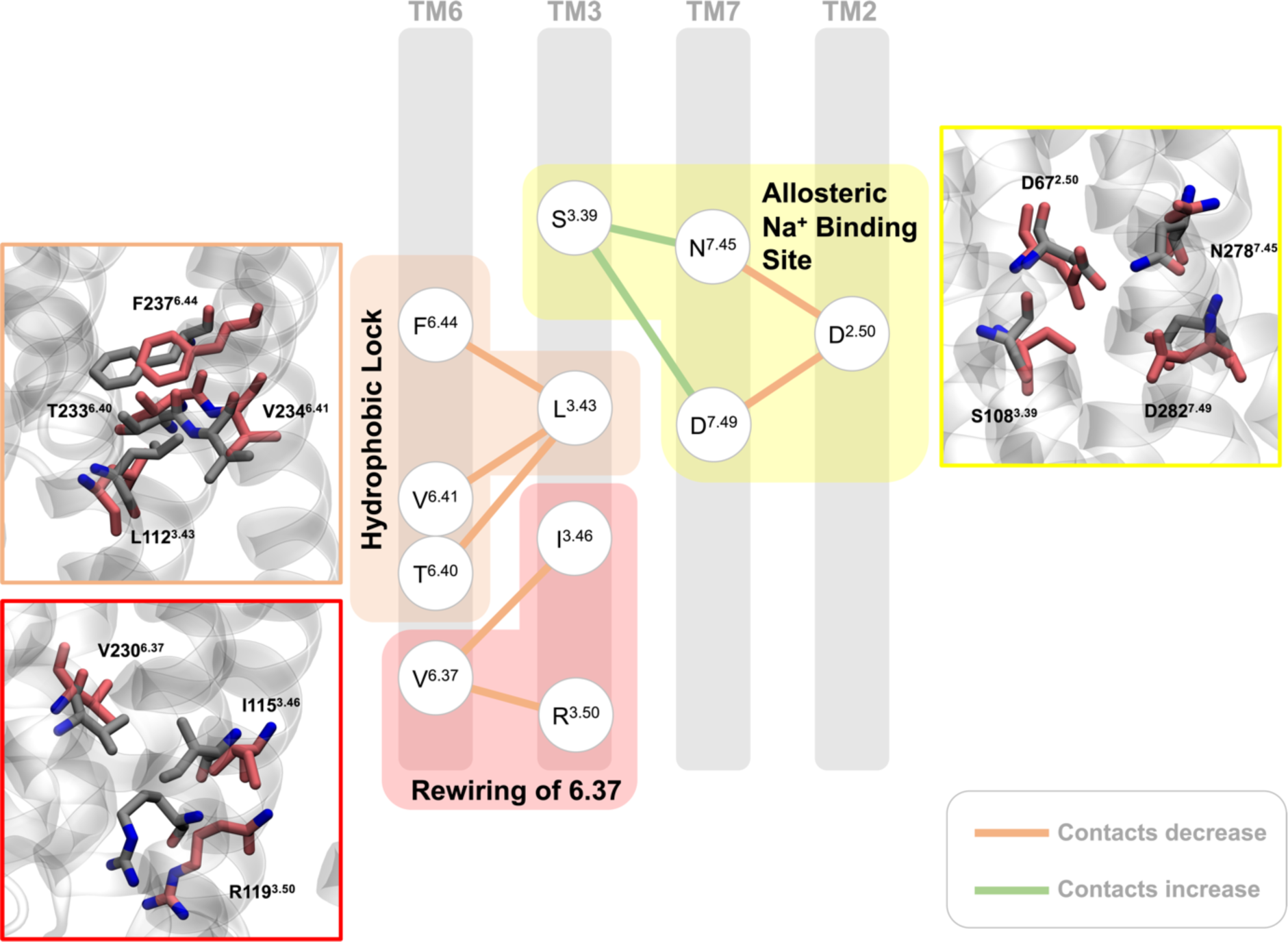
Microswitch changes in TM6 and TM7 in the GPR68 simulations. Schematic overview of changes in microswitches located in the region with largest motions along PC1. Microswitch residues are divided into three groups: the hydrophobic lock (orange), residue 6.37 (red) and the allosteric Na^+^ binding site (yellow). Residues are shown as nodes with the edges between them representing an increase (green) or a decrease (orange) of contacts upon activation determined by comparing their average distance in the ‘U’ inactive and active state replicas. Distances are calculated using the centre of mass of heavy atoms. For each group, a close-up view of representative snapshots of the MD1 pseudo-trajectory is also shown where the arrangement of microswitch residues at the beginning of the production run of the ‘U’ simulation (grey) and at the end of the production run of the ‘P6’ simulation (red) are shown as sticks.

The microswitch residue at position 6.37 (red spheres in Figure 5A and red shading in Figure 6.) is also found in the part of TM6 that shows a more pronounced motion in PC1. The change in its contacts is associated with the rearrangement of the intracellular side that eventually leads to G-protein binding^31^. During activation, a loss of contacts between this residue and TM3 residues at position 3.46 and 3.50 is usually observed. The V230^6.37^-R119^3.50^ and V230^6.37^- I115^3.46^ distances were monitored during the ‘U’ to ‘P6’ trajectories to detect any rewiring of 6.37 contacts upon protonation. Both distances were found to increase in MD1 and MD2 (Figure S8), indicating a rearrangement of these residues (red-lined close-up view in Figure 6.) and a transition towards the active state. As for the previous microswitch, a decrease or no significant change in these distances was observed for MD3.

At last, the behaviour of the residues lining the allosteric Na^+^ binding site in class A GPCRs (yellow spheres in Figure 5A and yellow shading in Figure 6.) was analysed. During activation, the inward movement of TM7 is expected to cause a rearrangement of these residues leading to a collapse of the binding site^31^. Monitoring distances involving residues D67^2.50^, S108^3.39^, N278^7.45^ and D282^7.49^ showed changes consistent with the system getting closer to the GPR68 active state in our MD1 and MD2 pseudo-trajectories (Figure S9). In MD3, the changes in these distances were found to be independent from what was expected for activation.

In summary, analysis of the inactive state simulations with increasing levels of protonation shows global (PC1) and local (microswitches) rearrangements consistent with a partial transition towards the active state for two out of three of our pseudo-trajectories.

### Stabilising hydrophobic interactions in the inactive state of GPR68

Our simulations indicate that protonation events on the extracellular side of GPR68 can propagate to the transmembrane and intracellular regions and initiate the transition towards the active state. To gain more insight into the residues involved in this mechanism, a Dynamic Cross-Correlation (DCC) analysis was performed on the pseudo-trajectories where global and local motions consistent with activation were observed (MD1 and MD2). A correlation network was generated on the basis of the DCC values and the shortest paths were determined that connect in the network each of the extracellular histidine residues protonated during the simulations (H17^1.28^, H84^2.67^ and H169^EL2^) to the three microswitches in the TM6 and TM7 regions. These shortest paths correspond to possible pathways of allosteric communication within the protein. The intermediate node residues in the paths were further analysed to identify residues that are consistently in contact with them either in the ‘U’ inactive or active state simulations. Contact pairs were then ranked according to their distances in the simulations and their conservation in the multiple sequence alignment of GPR68 homologs and a control alignment containing class A GPCRs with sequence identity to human GPR68 < 40%. Pairs were ranked higher if the distance between the residues in the pair differed significantly in the ‘U’ inactive and active simulations, if they were more conserved in GPR68 homologs and if they were less conserved in the control alignment (Supplementary Methods). The last contribution to the scoring function was added to identify pairs of residues involved in the activation mechanism that are unique to GPR68.

Mapping the top 20 ranked pairs (Figure S10) on the GPR68 structure (spheres in Figure 7A/B) shows that they are clustered in the transmembrane region closer to the extracellular side of the receptor, with the I15^1.26^ residue (paired with F265^7.32^) being the outermost of these residues and the closest to the three extracellular histidines considered in this analysis (green sticks). The top 20 pairs are mostly composed of hydrophobic residues (pink) and run from the extracellular side down to the microswitches (yellow, orange and red sticks) along TM1-3, TM6 and TM7.

**Figure 7.**
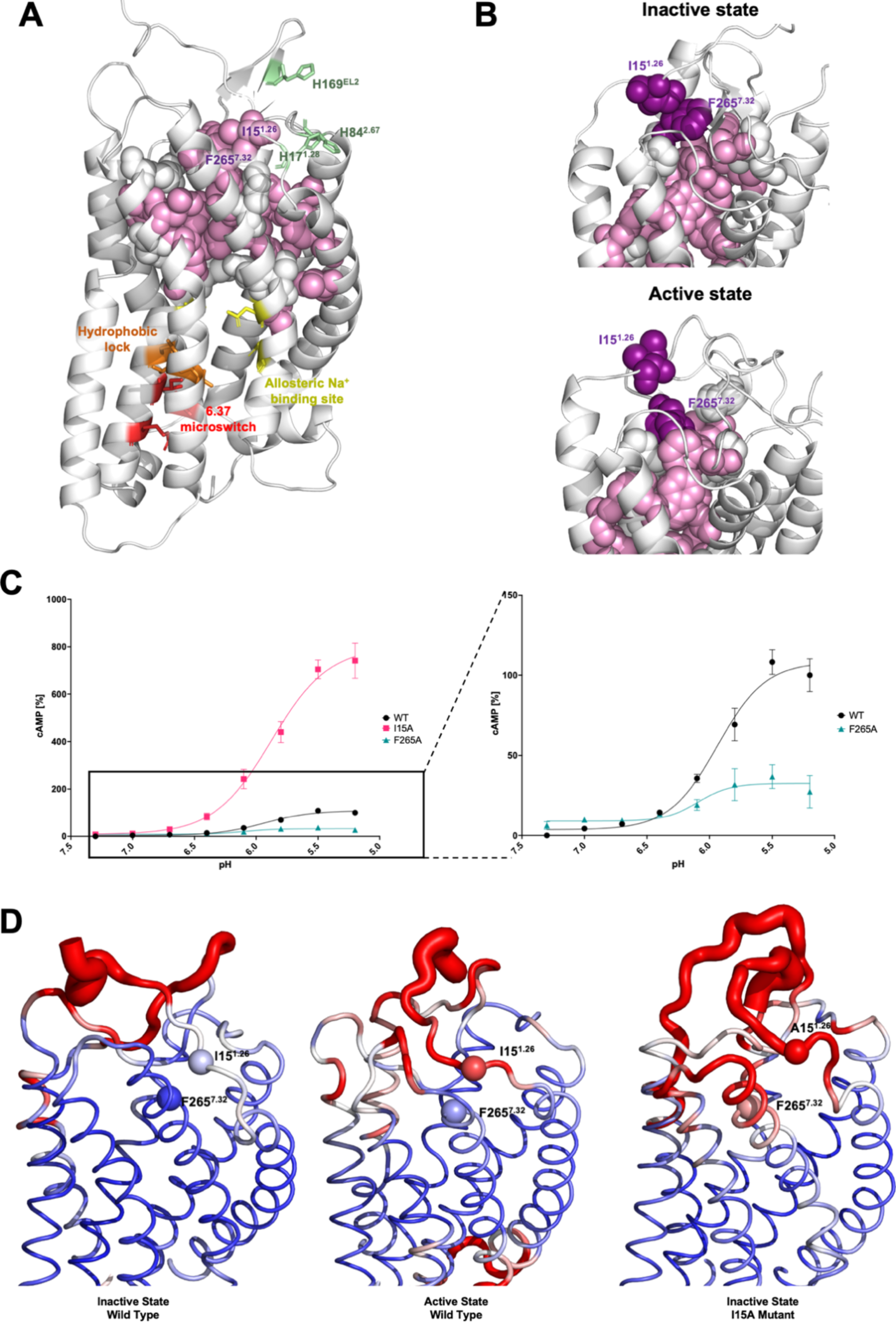
Extracellular hydrophobic interactions important for proton-sensing. A) The top-ranked residue pairs from the DCC analysis are shown in the representative structure of the ‘U’ inactive simulations (centroid of the most populated cluster). The top 20 pairs are shown as pink (hydrophobic residues) and white (polar/charged residues) spheres. The analysis to extract the pairs involved the calculation of the shortest paths connecting the three extracellular histidine residues H17^1.28^, H84^2.67^ and H169^EL2^ (green sticks) to three microswitch regions (yellow, orange and red sticks) in the DCC network. B) Close-up view of the I15^1.26^-F265^7.32^ pair (purple spheres) in the representative structure of the ‘U’ inactive (top) and active (bottom) state simulations. The other top 20 pairs are shown as spheres using the same colour code as in A. C) pH concentration-response curves for the cAMP production in HEK293 cells for the I15A and F265A mutants compared to wild type (left). A close-up view of the plot without I15A is shown on the right. Data represent the mean ± SEM from *n* = 3 individual replicates performed in triplicate. The error bars that are not visible lie within the dimensions of the symbol. D) Structural mapping of the Cα RMSF values for the wild type inactive (left) and active (middle) state, and the I15A mutant inactive state (right). RMSF values are colour-mapped from blue to red onto a representative structure of each state (centroid of the most populated cluster). The thickness of the tube representation is proportional to the RMSF value. Values were calculated using the production runs of all the replicas of the ‘U’ inactive, active and I15A mutant simulations.

The importance of the I15^1.26^-F265^7.32^ pair in the activation mechanism was further tested through mutagenesis studies. Remarkably, the I15A mutation (magenta in Figure 7C) was found to lead to an 8-fold gain of function compared to wild type (black), indicating that replacing the isoleucine in this position with a less bulky amino acid might destabilise the inactive state and favour the transition to the active one. Comparing the arrangement of the I15^1.26^-F265^7.32^ pair in the two states shows a loss of contact between these residues when going from the inactive to the active state (purple spheres in Figure 7B). Consistently, the Root Mean Square Fluctuation (RMSF) profiles calculated during the simulations show an increase in mobility in the region of the N-terminus around I15^1.26^ and of TM7 around F265^7.32^ in the active state compared to the inactive one (middle and left panel Figure 7D). A similar increase in flexibility was observed during the I15A mutant simulations starting from the inactive state (right panel), where the smaller alanine residue is not able to form the same interactions as the isoleucine and thus leads to a destabilisation of the inactive state arrangement in this region.

The other residue in the pair (F265^7.32^) is more buried than I15^1.26^ and, while it changes its arrangement when going from the inactive to the active state, it forms extensive contacts with surrounding hydrophobic residues in both states (Figure 7B). Correspondingly, mutating it to alanine produces a significant loss of function (cyan in Figure 7C).

In summary, we identified a group of hydrophobic residues in the region between proton-sensing histidines and known microswitches that undergo significant changes in their contacts upon activation, indicating they could be involved in propagating conformational changes from the extracellular side to the intracellular one. Both simulations and *in vitro* assays highlight the existence of a possible previously unknown hydrophobic lock formed by the I15^1.26^-F265^7.32^ pair, which regulates GPR68 activity by stabilising the inactive state conformation in the extracellular region.

## CONCLUSIONS

In this work, we investigated the activation mechanism of GPR68 using MD simulations and site-directed mutagenesis. A comprehensive modelling and simulation approach was adopted, where both the inactive and active states were studied following the same protocols. Starting from the inactive state, residues considered to be important for proton-sensing were sequentially protonated, with the sequence determined by the p*K*_a_ values of the residues observed during the simulation. Extensive analyses were performed to detect both local and global motions consistent with activation. New residues were identified as likely to be involved in the activation mechanism and validated through mutagenesis studies.

Among the extracellular histidine residues, H169^EL2^ was found to be the one more likely to be protonated at first on the basis of its predicted p*K*_a_ values, consistently with previous studies where H169^EL2^ and H20^1.31^ were shown to be more important than the others for proton activity^23^. In our simulations H20^1.31^ was never protonated as its p*K*_a_ values were always significantly downshifted, suggesting that the role of this residue in GPR68 activation might not be due to changes in its protonation state.

Our findings support the view that proton-sensing in GPR68 is not only due to histidine residues but acidic residues have a major role in it. Residue D67^2.50^ from the acidic triad consistently shows p*K*_a_ values over 6.8 for most of the time in all the simulations. Significant upward shifts in p*K*_a_ were also observed for the other triad residues (D282^7.49^ and E149^4.53^) in the later stages of the protonation sequence. Importantly, extracellular acidic residues D16^1.27^, D85^EL1^ and the triplet E164^EL2^ D165^EL2^ E166^EL2^ (EDE segment), while not directly changing their protonation state, were found to be paired with histidines H17^1.28^, H84^2.67^ and H169^EL2^ and to modulate their interactions during the simulations upon histidine protonation. The role of these acidic residues in activation was confirmed by our mutagenesis data which showed a drop in activity when these residues were mutated to alanine, with D85A showing the most prominent effect. Remarkably, D85^EL1^ has been suggested in the past to have a possible role in GPR68 regulation through interactions with H169^EL2^ on the basis of independently generated homology models^23^. Moreover, the acidic residues in our pairings have been recently found to be part of a water-mediated hydrogen bond network that might be involved in the activation of GPR68^24^.

The importance of the histidine-acidic residue pairing in the extracellular region of the receptor was further highlighted by our sequence analyses, which suggested that this type of pairing is also present in GPR4 and GPR65. Moreover, these pairings seem to go beyond H17^1.28^, H84^2.67^ and H169^EL2^ and include other histidine residues that, in the case of GPR68, have been shown to have a possible support role in proton-sensing (H89^EL1^, H159^4.63^, H175^EL2^)^23^. Sequential protonation of the acidic triad residues and extracellular histidines H17^1.28^, H84^2.67^ and H169^EL2^ triggered conformational changes consistent with the start of the transition towards the active state in two out of three sets of our replicas. These changes included concerted movements of the TM6 and TM7 helices and rearrangements of the hydrophobic lock, the microswitch residue 6.37 and the allosteric Na^+^ binding site that are consistent with activation^31^. Dynamic cross-correlation and contact analyses highlighted a cluster of hydrophobic residues potentially involved in propagating changes from the extracellular region to microswitches on the transmembrane side. Residues I15^1.26^ and F265^7.32^ were found to form a previously unknown hydrophobic lock that is open in the active state and closed in the inactive one, where the lock reduces the flexibility of the N-terminus and stabilises the inactive conformation in the extracellular region. These findings were corroborated by the *in vitro* experiments on the I15A mutant, which showed a remarkable 8-fold increase in activity compared to wild type.

This study significantly advances our understanding of the activation mechanism of GPR68 and other proton-sensing GPCRs. Our findings can be used in the future to guide the design of GPR68 modulators. Such compounds have been shown to have therapeutic potential *in vivo* in mouse models of colitis^33^ and the receptor itself is a promising target for the treatment of cancer^34^. GPR68 has been also found to be associated with genetic disorders^35^ and the identification of new mechanistically important amino acids may also help in the detection of further pathogenic variants.

## METHODS

### Molecular Dynamics Simulations

#### Generation of initial structures

For this work, the Ballesteros-Weinstein method of numbering residues was used^36^ (Table S4). Starting structures for the MD simulations of the inactive and active state of GPR68 were generated by homology modelling. Candidate templates were identified by using BLAST on the sequence of human GPR68 (UniProt ID: Q15743) against the Protein Data Bank database. The resulting structures were then mapped onto the GPCRdb database^25^ to extract their state annotation (active, inactive or intermediate) according to the delta distance *−d*:

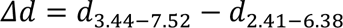

where *d_X-Y_* is the distance between the Cα atoms of residues X and Y^25^. Missing *Δd* values were calculated manually using VMD 1.9.4^37^. When more than one structure for a given receptor and state was present, only one was retained on the basis of the sequence identity to the target and the coverage. Structures with resolution worse than 3 Å were also removed.

The resulting templates were aligned using PRALINE^29^ with PSIPRED^38^ for secondary structure prediction and PHOBIUS^39^ for the prediction of transmembrane domains. For each state, multiple templates were selected (Tables S5 and S6) to improve the local match between template and target especially in the loop regions and termini. Similar approaches have been previously followed to model GPCR receptors^25^. A main template was first identified as the one best matching the transmembrane helices in terms of sequence identity and coverage. Secondary templates were then chosen for each loop and the termini of the receptor. The factors that influenced the selection were the coverage of the GPR68 sequence in each region, the similarity in sequence and the presence of polar amino acids and prolines. The homology models were generated using MODELLER 9.15^40^. A restraint needed to be added to model one of the expected^23^ GPR68 disulphide bonds (C13^1.24^-C258^EL3^) since this was not present in the templates. For the inactive model, an additional restraint of 4 Å with a 0.1 Å standard deviation was added between the Nε atoms of residues H17^1.28^ and H169^EL2^ to ensure proximity of these two residues and model the predicted extracellular cluster of histidine residues. These residues were chosen due to their position in regions of higher flexibility. For each state, a total of 1000 models was generated and the best model to be used as a starting point for the MD simulations was selected on the basis of the DOPE score^41^, match to existing experimental data and stability during preliminary MD simulations as detailed in the Supplementary Methods.

#### System setup and simulation protocol

The generated protein structures were embedded in a model membrane of 1-palmitoyl-2- oleoyl-sn-glycero-3-phosphocholine (POPC) lipids using the CHARMM-GUI Membrane builder^42,43^. The PPM 2.0 server^44^ was used to orient the protein in the membrane. A rectangular box was used, with water thickness (minimum height of water above and below the system) of 22.5 Å and an initial XY length of 80 Å, resulting in a number of POPC lipids for the lower and upper leaflets of ∼ 76-80.

MD simulations were performed with GROMACS 2020.3^45^. The system composed of the protein and the membrane was solvated with TIP3P molecules and counterions were added to neutralise it. Periodic boundary conditions were applied and a rectangular box was used. The protein, lipids and ions were described using CHARMM36m^46^. The Particle Mesh Ewald (PME) method^47^ was used for electrostatic interactions with a cutoff of 12 Å for direct-space sum. A 12-Å cutoff was also used for van der Waals interactions. The equations of motion were integrated using the leap-frog method with a 1-fs time step for most of the equilibration steps and 2-fs for the last equilibration step and the production run. LINCS^48^ was used to constrain bonds involving hydrogen atoms.

Each system was first energy-minimised using 50000-steps of steepest descent. Equilibration was carried out in different stages (Table S7). Positional restraints were first applied on both the protein and lipids heavy atoms and gradually decreased while equilibrating the temperature to 300K and the pressure to 1 bar. The Berendsen thermostat^49^ was used for the initial stage and subsequently replaced with Nosé-Hoover^50^. For the pressure coupling, the semi-isotropic Berendsen barostat^49^ was initially used and replaced by the Parrinello-Rahman^51^ for the last NPT stages. Production NPT runs were performed for 300 ns for all simulations. Three equilibration+production replicas were run for both the active and inactive states.

The starting structure of the I15A mutant trajectories was generated by using the mutagenesis wizard in PyMOL^52^ on the starting inactive structure. The same minimisation, equilibration and production protocol was used as described above.

#### Protonation protocol

Protonation events occurring within the GPR68 activation window were simulated by changing the ionisation state of residues on the basis of their p*K*_a_. Protonation was performed in stages and using p*K*_a_ values predicted with PROPKA 3.4.0^53^. An in-house Python code (available upon request) was used to run PROPKA on frames extracted every 100 ps from the inactive state simulations and containing both the protein and the lipid bilayer. The p*K*_a_ values of the different ionisable residues were compared to determine those most likely to change their protonation state within the GPR68 activation window and monitored during the simulations to identify frames to be protonated for the next stage.

When adding a proton to single residues, the frame with the highest p*K*_a_ in the last quarter of the production run was selected for protonation. When protonating two amino acids at the same time, the frame was selected so that it had the highest combined p*K*_a_ value in the last quarter of the production (as measured by the sum of the p*K*_a_ of each residue normalised as Z-score) and both residues had p*K*_a_ ≥ 6.8.

After each change in protonation, trajectories were started from the coordinates and velocities of the frame selected as described above to speed up equilibration. For the additional protons, initial coordinates were determined with PROPKA and null initial velocities were used. The overall neutrality of the system was maintained by replacing a water molecule with a Cl^-^ ion. Equilibration was run following steps 5-10 of the protocol described above (Table S7).

#### Trajectory analysis

Analyses were carried out on trajectory frames sampled every 100 ps unless otherwise stated. Representative structures for the inactive and active states were extracted by cluster analysis. For each state, clustering was performed using the *gromos* method^54^ on the concatenated production runs of the three replicas (‘U’ simulations were used for the inactive state). Structure dissimilarity was measured by calculating the Cα RMSD after removal of loop regions (residues 4-15, 50-53, 85-89, 126-133, 160-179, 215-218, 255-261, and, for the active state, 302-312). Optimal RMSD cutoff values were estimated to be 2.25 Å (inactive state and I15A mutant trajectories) and 2.50 Å (active state). The centroids of the most populated clusters were used as state representatives.

A Principal Component Analysis (PCA) was performed to identify one or more collective coordinates that could be used to monitor the inactive-to-active transition. The PCA was carried out with GROMACS on a pseudo-trajectory obtained by concatenating together the production phases of all the inactive state (‘U’ simulations) and active state replicas. Only the Cα atom coordinates in the transmembrane domain (residues 22-46, 59-80, 96-117, 137-158, 184-205, 229-249 and 264-284 from UniProt sequence annotation) were included in the pseudo-trajectory.

Dynamic cross-correlation (DCC) matrices were calculated for the inactive state replicas found to transition towards the active state upon protonation. Two pseudo-trajectories were first generated by concatenating the corresponding MD1 and MD2 replicas of the ‘U’ and ‘P2’ to ‘P6’ trajectories (Cα atoms only). The *dccm* function of the bio3d^55^ R library was applied to each pseudo-trajectory to generate two DCC matrices. A single consensus matrix was then determined, with elements set to the average of the elements from the individual matrices if they were both ≥ 0.4, and to 0 in all the other cases. The consensus matrix was used to define a correlation network with nodes (Cα atoms) joined by an edge if the corresponding element of the matrix is > 0. The shortest paths between groups of nodes in the network were calculated using the *cnapath* function from bio3d. Shortest paths were determined between each of the extracellular histidine residues selected for protonation (H17^1.28^, H84^2.67^ and H169^EL2^) and three microswitch regions in class A GPCRs (the hydrophobic lock (L112^3.43^, T233^6.40^, V234^6.41^ and F237^6.44^), residue 6.37 and nearby residues (V230^6.37^, I115^3.46^ and R119^3.50^) and the allosteric Na^+^ binding site (D67^2.50^, S108^3.39^, N278^7.45^ and D282^7.49^))^31^. The residues found in the paths were further filtered on the basis of a contact analysis and their evolutionary conservation as described in the Supplementary Methods.

Hydrogen bonds were detected with VMD version 1.9.4.^37^ using a donor-acceptor cutoff distance of 3.5 Å and a hydrogen-donor-acceptor angle of 30°.

### Sequence analysis of proton-sensing GPCRs

The sequences of human GPR68 (Uniprot ID: Q15743), GPR4 (Uniprot ID: P46093), GPR65 (Uniprot ID: Q8IYL9) and GPR132 (Uniprot ID: Q9UNW8) were run through BLAST^56^ against the non-redundant protein sequences database. For each receptor, a total of 5000 sequences were extracted and filtered to remove entries with words ‘like’, ‘low quality protein’, ‘hypothetical’ and ‘partial’ in the name. The resulting sequences were aligned using Fast M-Coffee^57^ and the sequence identity values from the alignment were used to remove sequences with identity to the original query < 40%. A refined multiple-sequence alignment was then generated by running Fast M-Coffee on the remaining sequences (Table S8).

For each histidine *i* in the human sequence of each receptor, the probability *P_i_* of finding an acidic residue near *i* in the sequence was calculated as the fraction of sequences for that receptor that have a D or E residue within two positions before or after histidine *i*. Only homologous sequences that have a histidine at position *i* in the multiple alignment were considered. The propensity of being in proximity of an acidic residue was then calculated as *P_i_/P_ave_*, where *P_ave_*is the average probability calculated over all the histidine residues of the human sequences. In the subsequent analyses, the histidine residues were split into different groups. Their classification into extracellular, transmembrane, or intracellular was guided by the feature section of the human sequences from UniProt. The extracellular histidine residues were further classified as “involved” or “not involved” in proton-sensing based on experimental data from published mutagenesis experiments (Table S2). If no experimental data could be found, the extracellular histidine residues were classified as “not tested”.

### *In vitro* experiments

#### Transient Transfection

Reverse transfection was used to transiently transfect HEK293 cells using Lipofectamine 3000 (Thermo Fisher), using the method provided by the manufacturer. Transfections were performed such that each well contained 100 ng of the GPR68 receptor (wild type or mutant, Twist Biosciences) and 50 ng of the pGlo-Sensor^TM^-22F cAMP protein sensor with lipofectamine added in a 1:1.5 w:v ratio, respectively. A total of 50 μL of this mix was added to a poly-D-lysine (Sigma-Aldrich) coated F white clear-bottom plate (Greiner Bio-One). To this, 100 μL of HEK293 cells at a viable cell density of 75 000 cells was then added. Plates were then incubated in a 5% CO_2_ atmosphere at 37 °C for 24 h prior to performing intracellular cAMP production assays.

#### Intracellular cAMP Assays

Twenty-four hours post-transfection, the cell culture media were removed slowly, minimising disruption to attached cells adhered to the bottom of the wells. Cells were initially washed using HBSS-based cAMP assay buffer (pH 7.3). Thereafter, cAMP buffer supplemented with firefly D-luciferin (0.45 mg/mL; NanoLight Technologies) was added (100 μL). The plate was then left to pre-equilibrate in the dark at 28 °C for 1 h. During this time, the CLARIOstar PLUS (BMG Labtech) was set to 28 °C. Aliquots of the buffer were taken, and the pH was adjusted using NaOH or HCl at 1 M. A total of 8 buffers were prepared with pH values of 7.3, 7.0, 6.7, 6.4, 6.1, 5.8, 5.5, and 5.2. Bioluminescence readings were then conducted to measure basal luminescence signal (5 cycles) prior to buffer addition. After the baseline was read, 90 μL of the solution were removed from each well, and 90 μL of the necessary pH-adjusted buffer were added. Upon acidification through the buffer, luminescence readings were taken for 30 min. The area under the curve of the readings was used as a measurement. Each time point was normalised to the mean baseline reading of the equivalent well. GraphPad Prism 9.0 was used to analyse and plot data. It is to be noted that here GPR68 activity is measured as cAMP production instead of cAMP accumulation, which leads to a shift in our activation window of GPR68 compared to values previously reported in the literature^23^.

## ACKNOWLEDGEMENTS

We would like to thank Dr Alessandro Pandini from Brunel University for insightful discussions about the sequence analyses presented in this work.

This work was supported by the Biotechnology and Biological Sciences Research Council [grant number BB/M009513/1] and made use of time on HPC granted via the UK High-End Computing Consortium for Biomolecular Simulation, HECBioSim (http://hecbiosim.ac.uk), supported by EPSRC (grant no. EP/R029407/1).

