## Supplementary Information for "Elucidating the activation mechanism of the proton-sensing GPR68 receptor"

### Supplementary Methods

#### Molecular Dynamics Simulations

*Generation of initial structures - Inactive state*

The best inactive model among the 1000 models generated by MODELLER[1] was selected on the basis of the following pipeline.

The models were first ranked on the basis of their DOPE score[2] and of the distances between selected histidine residues predicted in previous work to form a cluster[3] ($H17^{1.28}$, $H20^{1.31}$, $H84^{2.67}$, $H169^{EL2}$ and $H269^{7.36}$). Distances were calculated between the Nε nitrogen atoms of each pair of residues and they were summed together to obtain the sum of distances to $H17^{1.28}$ (due to the flexibility of the N-terminus and the restraints added between $H17^{1.28}$ and $H169^{EL2}$ in the generation of the models, $H17^{1.28}$ was chosen as the centre of the cluster) and the sum of the distances between all pairs of histidines. The overall score used for the ranking was calculated as a combination of the normalised distance sums and DOPE score, where normalisation was carried out by calculating the corresponding Z-scores. Models with lower overall score were ranked higher and the top 20 models were selected as the models with the best combination of DOPE score and distances (with lower distances more likely to correspond to histidine clusters).

The selected models were further filtered on the basis of their match to literature information on antagonists. Two compounds (2-((2-ethyl-5,7-dimethyl-3H-imidazo[4,5-b]pyridin-3-yl)methyl)-8-((4-methylpiperazin-1-yl)methyl)-10,11-dihydro-5H-dibenzo[b,f]azepine and 2-((2-ethyl-5,7-dimethyl-3H-imidazo[4,5-b]pyridin-3-yl)methyl)-8-(piperidin-1-ylmethyl)-10,11-dihydro-5H-dibenzo[b,f]azepine) known to have a different degree of antagonistic activity on GPR68[4] and differing only by one methyl group were docked to each of the top 20 models. For each model, six candidate binding sites known from the literature[5] were considered, with docking calculations performed separately for each binding site using Autodock Vina[6] and grid boxes defined to fit the amino acids in the site. The exhaustiveness was set to 100. The binding sites and the models were ranked on the basis of their ability to explain the difference in activity between the two compounds and the top 8 models were retained for the next stage.

Molecular Dynamics (MD) simulations were run on the selected models, following the same protocol as described in the main text, with production runs of 300 ns. The stability of receptor during the MD simulations was monitored through calculation of the Root Mean Square Deviation (RMSD) of the Cα atoms from the initial structure. The top three models were selected based on their stability during MD and of the proximity of histidine residues on the extracellular side.

To take into account changes in the shape of the binding pocket observed during the MD simulations, the two antagonists were re-docked to representative structures extracted from the trajectories. A cluster analysis was performed with the *gromos* clustering method[7] implemented in GROMACS[8] on structures sampled every 100 ps using a cutoff of 3.5 Å. A total of 13 cluster representatives were extracted from the three trajectories. The RMSD values used for the clustering were calculated on the non-hydrogen atoms of the binding site residues (determined as within 4 Å of the docked ligand in the three initial models). MD simulations were run on selected docked structures showing recurrent binding poses. The ligands were described with the CGenFF force field[9]. The model with the highest stability during these simulations was finally selected to study the activation mechanism of GPR68.

*Generation of initial structures - Active state*

The active state model was chosen based on its consistency to literature data on binding of ogerin[3] and other Positive Allosteric Modulators (PAM). The top 20 models based on the

DOPE score were selected from the initial 1000 models generated by Modeller and ogerin was docked to its predicted[3] binding site using Autodock Vina. A 14x14x14 Å$^3$ grid box was defined to include the binding site and exhaustiveness was set to 1000. A total of nine binding poses were generated for each model and the minimum distance of ogerin to the mutagenesis-validated binding site residues E160$^{4.64}$, R189$^{5.42}$ and H269$^{7.36}$ was determined for each pose (heavy atoms only). Poses were ranked on the basis of a scoring function calculated as a combination of the normalised distances and binding affinity, where normalisation was carried out by calculating the corresponding Z-scores. The top 9 models that most frequently showed binding poses with low Z-scores were selected.

A library of known GPR68 PAMs (92 compounds)[3,10] was docked to each selected model, using the same docking box as ogerin and an exhaustiveness of 500. Models were ranked according to the agreement of the corresponding docking results with available allosteric parameters. The two best models were retained for MD simulations, and the one showing the highest stability was selected as the final active model.

*Contact analysis and ranking of residue pairs*

A contact analysis was performed on the residues making up the shortest paths in the DCC network between selected extracellular histidine residues and microswitches described in the main text. The contact frequency between each residue in the paths and the rest of the protein during the 'U' inactive and active simulations was calculated using the Contact Map Explorer[11] library for Python. A contact between a residue pair was identified when the distance between any atoms in their side chains (heavy atoms only) was found below 4.5 Å. Only the pairs with a contact frequency > 0 in all the replicas for at least one state (inactive or active) were retained for further analyses.

The selected pairs were classified into four groups on the basis of the distribution of their distances (calculated as minimum distances between the side chain heavy atoms) in the 'U' inactive and active state simulations. Pairs were assigned to the 'no change' group if their inactive and active distance distributions did not show a statistically significant difference as determined by a t-test ($p>0.05$), the 'always in contact' group if the mean for both distributions was ≤ 4.5 Å, the 'no contact' group if the mean of each distribution was > 4.5 Å and the 'change in contact' group if the mean of only one distribution was ≤ 4.5 Å. Only the pairs in the last group were selected for the next step and the dissimilarity between their inactive and active state distance distributions was quantified by calculating their Bhattacharyya distance with the *dictances* Python library (version 2.5.6, https://pypi.org/project/dictances/).

Conservation scores were calculated for each residue in the selected pairs using the multiple sequence alignment of GPR68 homologues described in the main text and a control alignment containing human class A GPCR sequences. The latter was generated by running the sequence of human GPR68 (Uniprot ID: Q15743) through BLAST[12] against the non-redundant protein sequences database and applying filters to only include results from *Homo sapiens*. A total of 859 sequences were extracted and filtered to remove entries with words 'like', 'low quality protein', 'hypothetical' and 'partial' in the name. Entries with an E-value > 0.01 (max E-value = 0.046) were further examined to confirm they are class A GPCRs. The sequences were aligned using Fast M-Coffee[13] and the sequence identity values from the alignment were used to remove sequences with identity to the original query > 40%. A refined multiple-sequence alignment was then generated by running Fast M-Coffee on the remaining sequences and used as control alignment. Conservation scores were calculated for both alignments using the similarity method (bio3d similarity matrix) of the *conserv* function from the R library bio3d[14].

For each pair of amino acids $X$ and $Y$, the Z-score of the Bhattacharyya distance ($Z_{Bh}$) was calculated. The conservation score in the GPR68 homolog alignment ($Z_{GPR68}$) and in the control alignment ($Z_C$) was also calculated for each amino acid. The pairs were ranked according to the linear combination *Z*:

$$Z = Z_{Bh} + Z_{X,GPR68} + Z_{Y,GPR68} - Z_{X,C} - Z_{Y,C}$$

**Table S1.** Trajectory data for the simulations with different degrees of protonation of selected histidine and acidic residues. The frame selected for residue protonation is indicated for every replica (MD1, MD2 and MD3).

| Trajectory | Protonation | Duration [ns] | MD1 [ps] | MD2 [ps] | MD3 [ps] |
|---|---|---|---|---|---|
| U | None[a] | 300 | 281500 | 240200 | 202300 |
| P2 | $D67^{2.50}$, $H169^{EL2}$ | 300 | 219400 | 207100 | 262200 |
| P3 | $H17^{1.28}$, $D67^{2.50}$, $H169^{EL2}$ | 300 | 206600 | 210900 | 252300 |
| P4 | $H17^{1.28}$, $D67^{2.50}$, $H84^{2.67}$, $H169^{EL2}$ | 900 | 664200 | 844390 | 656300 |
| P5 | $H17^{1.28}$, $D67^{2.50}$, $H84^{2.67}$, $H169^{EL2}$, $D282^{7.49}$ | 600 | 470200 | 431300 | 436900 |
| P6 | $H17^{1.28}$, $D67^{2.50}$, $H84^{2.67}$, $E149^{4.53}$, $H169^{EL2}$, $D282^{7.49}$ | 600 | - | - | - |

[a]Ionisable residues set to their standard protonation state at pH 7

**Table S2** Classification of histidine residues in GPR68, GPR4, GPR65 and GPR132. All histidine residues are categorised according to their location in the receptor and mutagenesis data from the literature.

| **Receptor**[a] | **Extracellular – involved in proton-sensing** | **Extracellular – not involved in proton-sensing** | **Extracellular – role in proton sensing not experimentally tested** | **Transmembrane** | **Intracellular** |
|---|---|---|---|---|---|
| GPR68[15,16] | H17$^{1.28}$, H20$^{1.31}$, H84$^{2.67}$, H169$^{EL2}$ (main) H89$^{EL1}$, H159$^{4.63}$, H175$^{EL2}$ (cooperative effect with main residues) | - | - | H245$^{6.52}$, H269$^{7.36}$ | H125$^{3.56}$, H130$^{IL2}$, H216$^{IL3}$, H294$^{8.51}$, H345$^{CTER}$ |
| GPR4[16,17] | H79$^{2.66}$, H165$^{EL2}$ | H4$^{NTER}$, H10$^{NTER}$, H17$^{1.32}$, H80$^{2.67}$, H85$^{EL1}$ | H155$^{4.63}$ | H241$^{6.52}$, H269$^{7.36}$ | H121$^{3.56}$, H302$^{8.59}$ |
| GPR65[16,18] | H10$^{NTER}$, H14$^{1.32}$ | - | H254$^{6.63}$, H261$^{7.24}$ | H243$^{6.52}$ | H212$^{5.67}$ |
| GPR132[19] | - | - | H106$^{EL1}$ | H174$^{4.57}$, H220$^{5.59}$, H259$^{6.52}$ | H242$^{6.35}$, H314$^{8.49}$, H323$^{8.58}$, H340$^{CTER}$, H356$^{CTER}$, H364$^{CTER}$ |

[a] References containing the data used for the classification of histidine residues are cited for each receptor

**Table S3.** Variance and cumulative variance of the first five principal components (PC) calculated for the pseudo-trajectory composed of the 'U' inactive state and active state replicas.

| PC | Variance [%] | Cumulative variance [%] |
|---|---|---|
| 1 | 48.2 | 48.2 |
| 2 | 15.2 | 63.4 |
| 3 | 10.3 | 73.7 |
| 4 | 7.4 | 81.1 |
| 5 | 3.4 | 84.5 |

**Table S4.** Ballesteros and Weinstein (BW) numbering[20] for the GPR68, GPR4, GPR65 and GPR132 residues mentioned in this work.

| BW numbering | GPR68 | GPR4 | GPR65 | GPR132 |
|---|---|---|---|---|
| NTER | - | H4, H10 | H10 | - |
| 1.24 | C13 | - | - | - |
| 1.26 | I15 | - | - | - |
| 1.27 | D16 | - | - | - |
| 1.28 | H17 | - | - | - |
| 1.31 | H20 | - | - | - |
| 1.32 | - | H17 | H14 | - |
| 1.39 | Y28 | - | - | - |
| 1.42 | V31 | - | - | - |
| 2.38 | E55 | - | - | - |
| 2.50 | D67 | - | - | - |
| 2.53 | Y70 | - | - | - |
| 2.60 | W77 | - | - | - |
| 2.66 | - | H79 | - | - |
| 2.67 | H84 | H80 | - | - |
| EL1 | D85, H89 | H85 | - | H106 |
| 3.22 | D91 | - | - | - |
| 3.32 | L101 | - | - | - |
| 3.34 | E103 | - | - | - |
| 3.35 | N104 | - | - | - |
| 3.36 | I105 | - | - | - |
| 3.39 | S108 | - | - | - |
| 3.49 | D118 | - | - | - |
| 3.56 | H125 | H121 | - | - |
| IL2 | H130 | - | - | - |
| 4.53 | E149 | - | - | - |
| 4.57 | - | - | - | H174 |
| 4.63 | H159 | H155 | - | - |
| 4.64 | E160 | - | - | - |
| EL2 | E161, E164, D165, E166, H169, F173, E174, H175 | H165 | - | - |
| 5.59 | - | - | - | H220 |

| | | | | |
|---|---|---|---|---|
| 5.67 | - | - | H212 | - |
| IL3 | H216 | - | - | - |
| 6.31 | D224 | - | - | - |
| 6.35 | - | - | - | H242 |
| 6.51 | Y244 | - | - | - |
| 6.52 | H245 | H241 | H243 | H259 |
| 6.54 | L247 | - | - | - |
| 6.55 | L248 | - | - | - |
| 6.58 | R251 | - | - | - |
| 6.61 | W254 | - | - | - |
| 6.63 | - | - | H254 | - |
| EL3 | E255, C258, D259 | - | - | - |
| 7.24 | - | - | H261 | - |
| 7.28 | A261 | - | - | - |
| 7.32 | F265 | - | - | - |
| 7.35 | Y268 | - | - | - |
| 7.36 | H269 | H269 | - | - |
| 7.39 | L272 | - | - | - |
| 7.42 | T275 | - | - | - |
| 7.43 | S276 | - | - | - |
| 7.46 | C279 | - | - | - |
| 7.49 | D282 | - | - | - |
| 8.48 | E291 | - | - | - |
| 8.49 | - | - | - | H314 |
| 8.51 | H294 | - | - | - |
| 8.53 | D296 | - | - | - |
| 8.58 | - | - | - | H323 |
| 8.59 | - | H302 | - | - |
| CTER | H345 | - | - | H340, H356, H364 |

**Table S5.** Templates selected for the homology models of the active and inactive state of GPR68. Structures used are indicated with their PDB ID.

| **State** | Inactive | Active |
|---|---|---|
| **Residue Interval** | I4-R312 | I4-R301 |
| **Main template** | 5O9H | 6DO1 |
| **N-terminus (I4$^{NTER}$-C13$^{1.24}$)** | 6DO1 | 6DO1 |
| **IL1 (Q49$^{1.60}$-N54$^{2.37}$)** | 5O9H | 5XSZ |
| **EL1 (H84$^{2.67}$-G90$^{3.21}$)** | 5O9H | 6RNK |
| **IL2 (H125$^{3.56}$-T134$^{4.38}$)** | 4N6H | 5C1M |
| **EL2 (E160$^{4.64}$-P177$^{5.30}$)** | 5O9H | 6DO1 |
| **IL3 (R214$^{5.67}$-Q219$^{6.26}$)** | 5LWE | 5UNF |
| **EL3 (W254$^{6.61}$-A261$^{7.28}$)** | 4ZUD | 6RNK |
| **C-terminus (inactive: V289$^{7.56}$-R312$^{CTER}$) (active: V289$^{7.56}$-R301$^{8.58}$)** | 5O9H | 6DO1 |

**Table S6.** Details of selected templates.

| UniProt ID | Receptor family | Species | PDB ID | Resolution [Å] | State | E value | Sequence identity [%] | Coverage [%] |
|---|---|---|---|---|---|---|---|---|
| AGTR1 | Angiotensin | Human | 6DO1 | 2.9 | Active | 3.00E-27 | 31.68 | 54 |
| Q08BG4 | Lysophospholipid (LPA) | Zebrafish | 5XSZ | 3.2 | Active | 1.00E-27 | 32.69 | 55 |
| Q6IYF9 | Succinate | Rat | 6RNK | 1.94 | Active | 6.00E-23 | 27.15 | 78 |
| OPRM | Opioid | Mouse | 5C1M | 2.1 | Active | 1.00E-24 | 27.44 | 73 |
| AGTR2 | Angiotensin | Human | 5UNF | 2.8 | Active | 5.00E-24 | 28.47 | 76 |
| C5AR1 | Complement peptide | Human | 5O9H | 2.7 | Inactive | 6.00E-20 | 27.21 | 73 |
| OPRD | Opioid | Human | 4N6H | 1.8 | Inactive | 2.00E-19 | 28.47 | 73 |
| CCR9 | Chemokine | Human | 5LWE | 2.8 | Inactive | 1.00E-24 | 25.61 | 75 |
| AGTR1 | Angiotensin | Human | 4ZUD | 2.8 | Inactive | 9.00E-36 | 29.08 | 74 |

**Table S5.** Sequence of equilibration steps for the MD simulations.

| Step | Duration [ns] | Conditions | Restraints on protein heavy atoms [kJ/mol/nm$^2$] | Restraints on lipid heavy atoms [kJ/mol/nm$^2$] |
|---|---|---|---|---|
| 1 | 1 | NVT | 1000 | 1000 |
| 2 | 5 | NPT | 1000 | 1000 |
| 3 | 5 | NPT | 1000 | 500 |
| 4 | 5 | NPT | 1000 | 250 |
| 5 | 10 | NPT | 1000 | - |
| 6 | 5 | NPT | 500 | - |
| 7 | 5 | NPT | 500 (only Cα atoms) | - |
| 8 | 5 | NPT | 250 (only Cα atoms) | - |
| 9 | 20 | NPT | - | - |
| 10 | 30 | NPT | - | - |

**Table S8**. Number of sequences and maximum E-value in the multiple sequence alignments for proton-sensing receptors GPR4, GPR65, GPR68 and GPR132

| **Receptor** | **Number of sequences** | **Maximum E-value** |
|---|---|---|
| GPR68 | 702 | 9e-109 |
| GPR4 | 365 | 1e-64 |
| GPR65 | 613 | 1.2e-91 |
| GPR132 | 278 | 6e-106 |

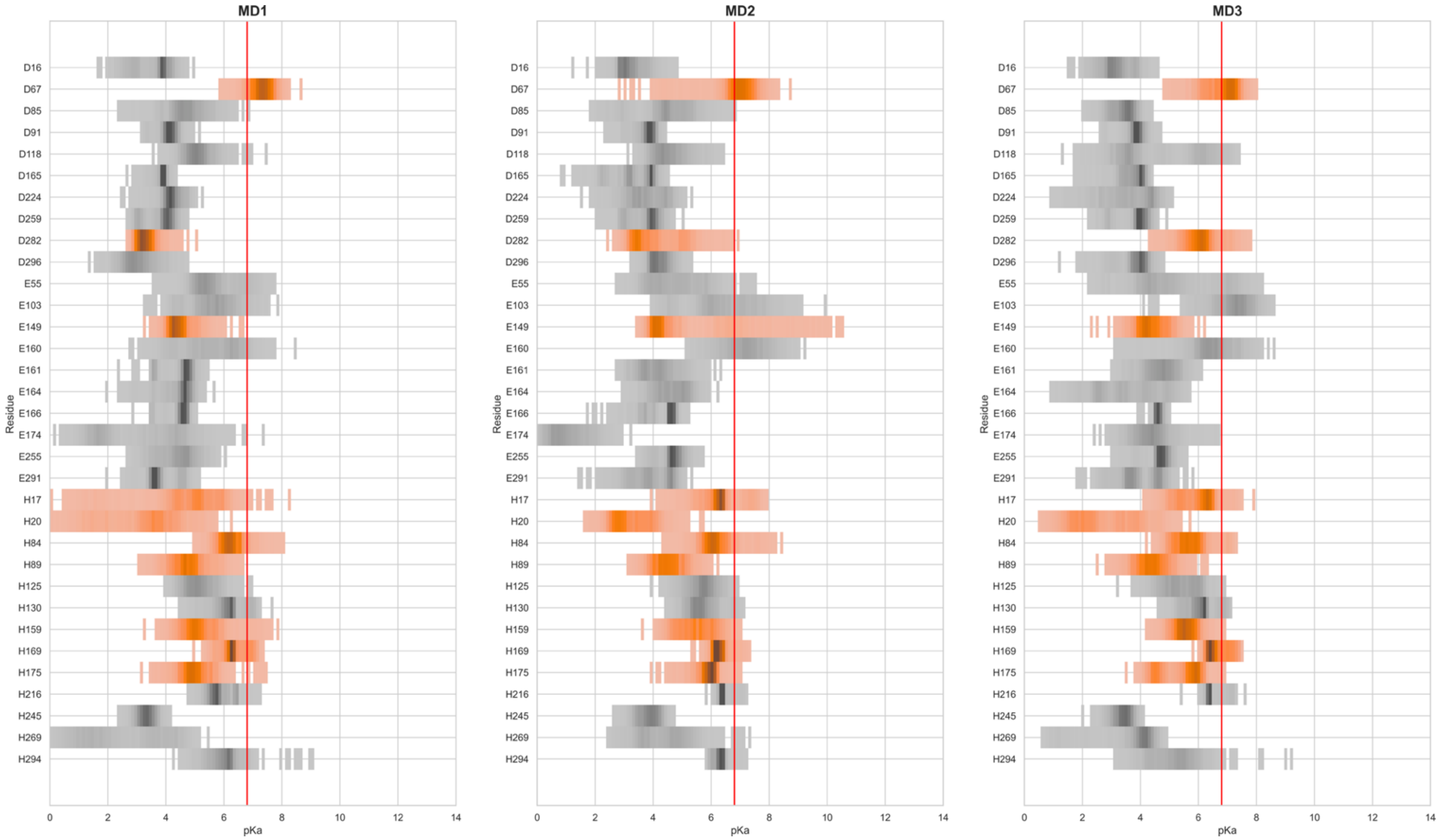


**Figure S1.** Distribution of predicted p$K_a$ values for acidic and histidine amino acids in GPR68. Values were calculated on the 300-ns production portion of the three 'U' replicas. A red vertical line is drawn at 6.8 (proton $EC_{50}$ value for GPR68[15]). The residues forming the acidic triad and the extracellular histidine residues are highlighted in orange.

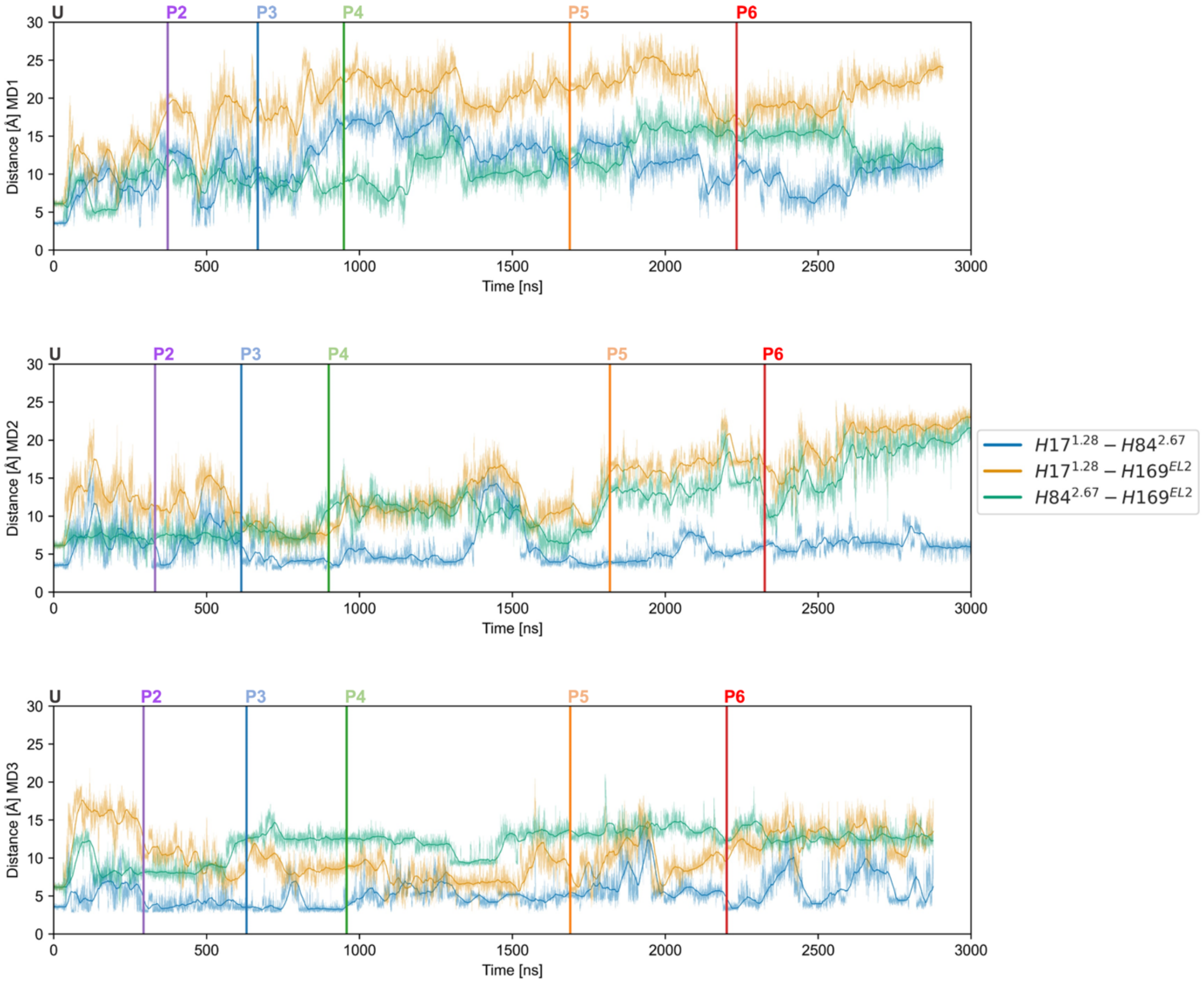


**Figure S2.** Time evolution of distances between histidine residues during MD simulations. The plots show the running average (30-ns window) of the minimum distances between $H17^{1.28}$-$H84^{2.67}$ (blue), $H17^{1.28}$-$H169^{EL2}$ (yellow) and $H84^{2.67}$-$H169^{EL2}$ (green) during the MD1 (top), MD2 (middle) and MD3 (bottom) simulations. Minimum distances are calculated over all possible pairs of heavy atoms from the two residues. Vertical lines separate the individual trajectories with different protonation states (‘U’ to ‘P6’).

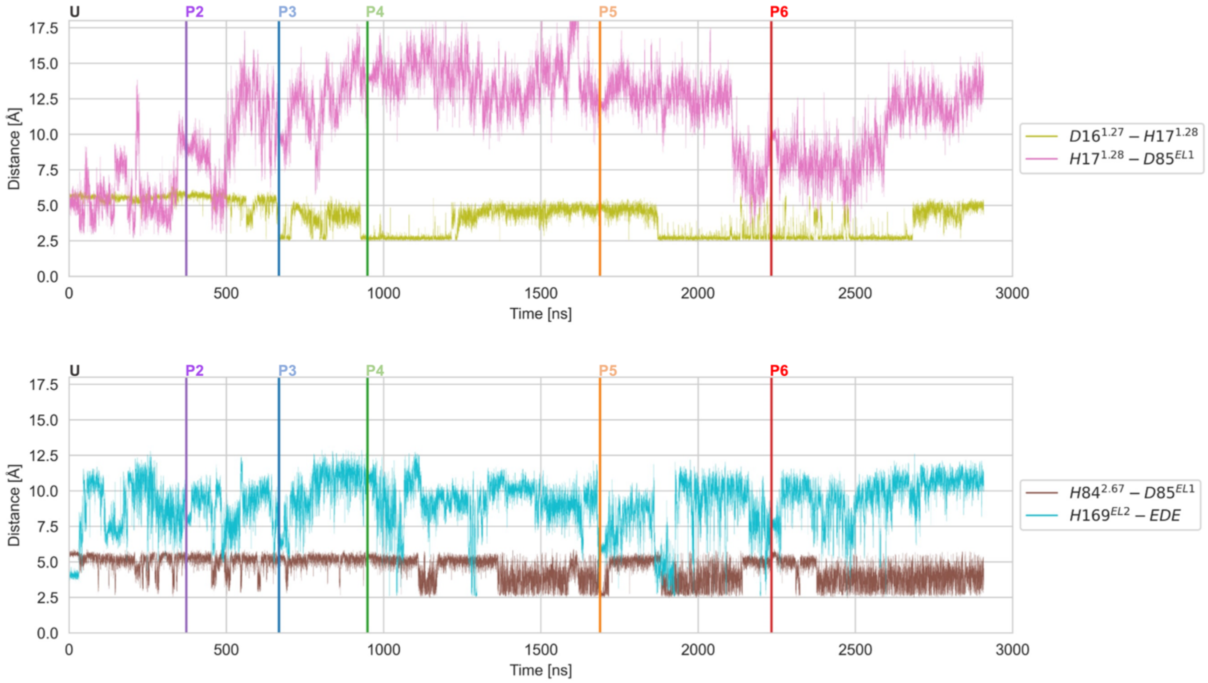


**Figure S3.** Time evolution of selected distances between histidine and acidic residues during the MD1 pseudo-trajectory. The minimum distance calculated over all possible pairs of side chain heavy atoms is plotted for the D16$^{1.27}$-H17$^{1.28}$ and H17$^{1.28}$-D85$^{EL1}$ (top panel) and H84$^{2.67}$-D85$^{EL1}$ and H169$^{EL2}$-EDE (bottom panel) pairs. Vertical lines separate the individual trajectories with different protonation states (‘U’ to ‘P6’).

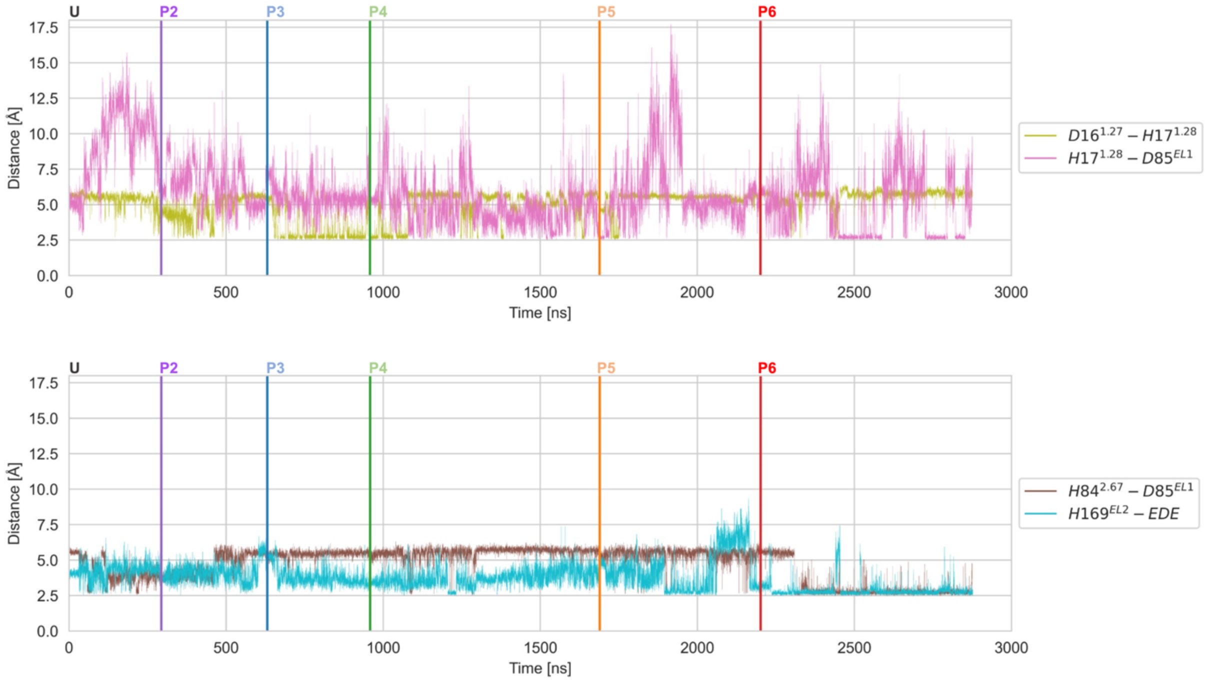


**Figure S4.** Time evolution of selected distances between histidine and acidic residues during the MD3 pseudo-trajectory. The minimum distance calculated over all possible pairs of side chain heavy atoms is plotted for the $D16^{1.27}$-$H17^{1.28}$ and $H17^{1.28}$-$D85^{EL1}$ (top panel) and $H84^{2.67}$-$D85^{EL1}$ and $H169^{EL2}$-EDE (bottom panel) pairs. Vertical lines separate the individual trajectories with different protonation states (‘U’ to ‘P6’).

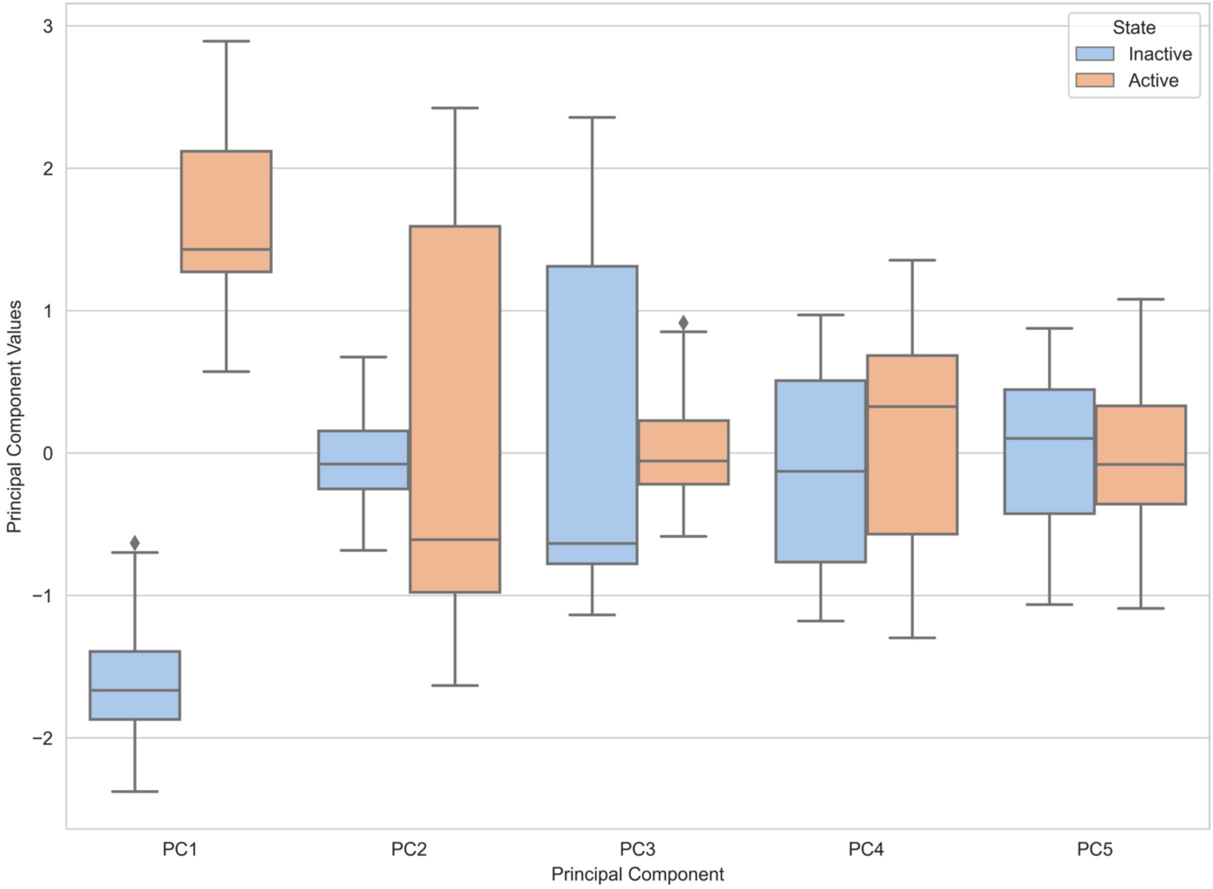


**Figure S5.** Projection of the inactive and active state trajectories on the first five principal components. Values calculated on the production runs of the 'U' inactive and active state replicas are shown as boxplots. The box indicates the central quartiles of the distribution separated by the median (vertical line). The whiskers show the rest of the distribution except for the outliers (diamond shapes).

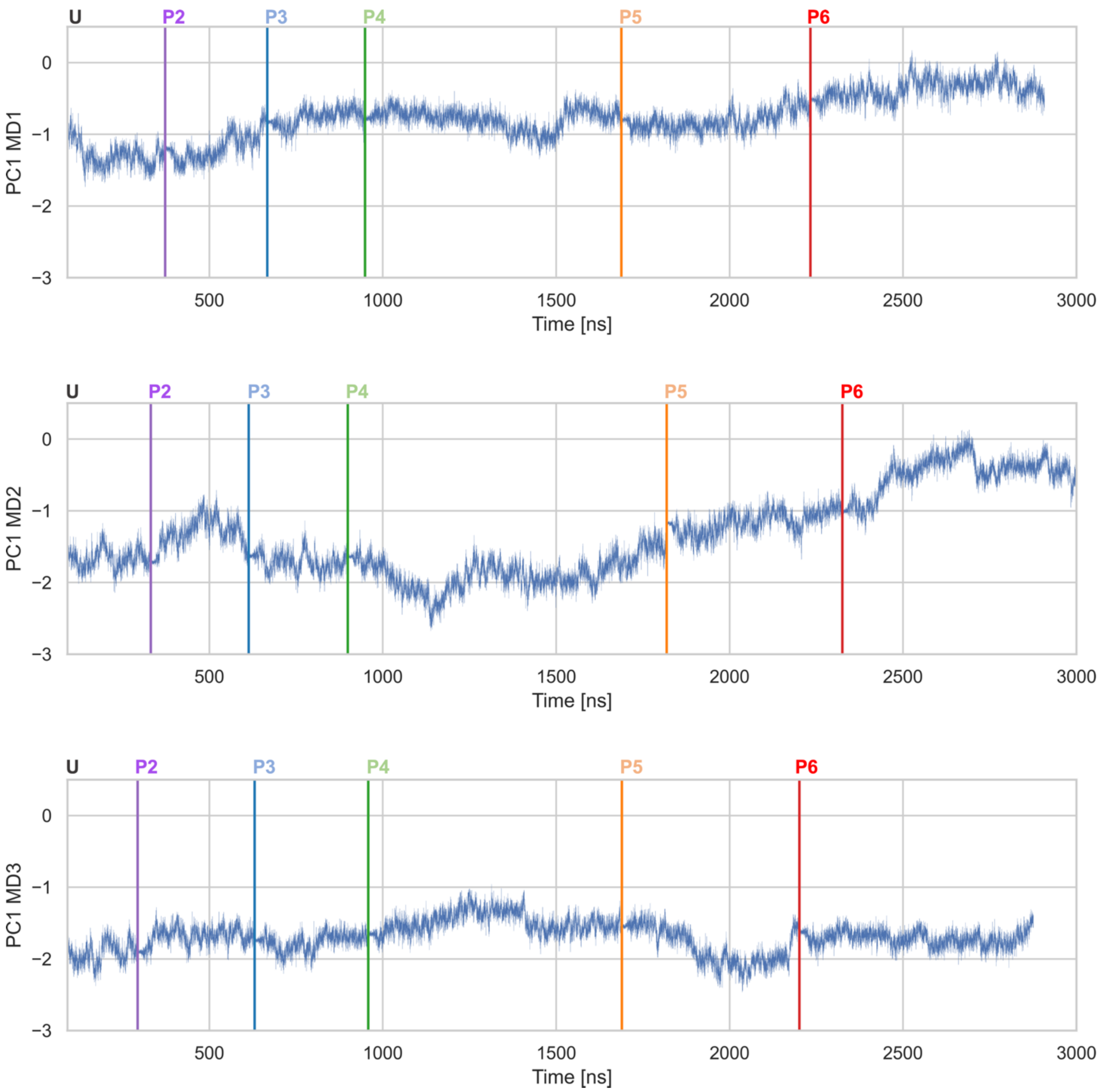


**Figure S6.** Time evolution of the projection of the MD1, MD2 and MD3 pseudo-trajectories on PC1. Vertical lines separate the individual trajectories with different protonation states ('U' to 'P6' simulations). The initial equilibration of the 'U' trajectories is not shown.

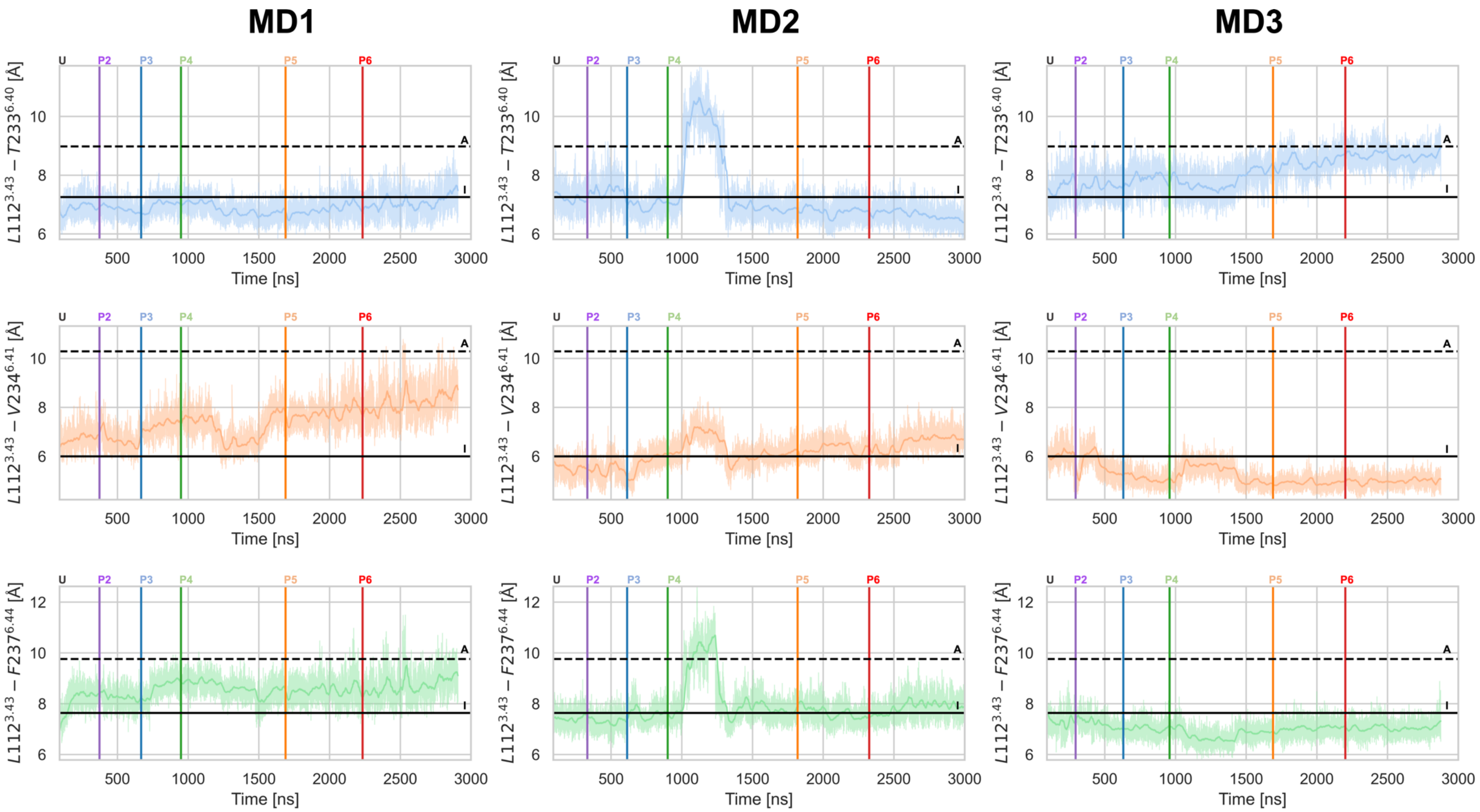


**Figure S7.** Time evolution of hydrophobic lock distances. The L112$^{3.43}$-T233$^{6.40}$ (blue), L112$^{3.43}$-V234$^{6.41}$ (orange), and L112$^{3.43}$-F237$^{6.44}$ (green) distances are shown for the MD1 (left), MD2 (middle) and MD3 (right) pseudo-trajectories. Distances were calculated between the centres of mass of the heavy atoms for each residue pair. Running averages (30-ns window) are shown with a darker hue. Vertical lines separate the individual trajectories with different protonation states (‘U’ to ‘P6’ simulations). The average distance values calculated over all the ‘U’ inactive (solid black line) and the active (dashed black line) state replicas (production only) are shown as horizontal lines.

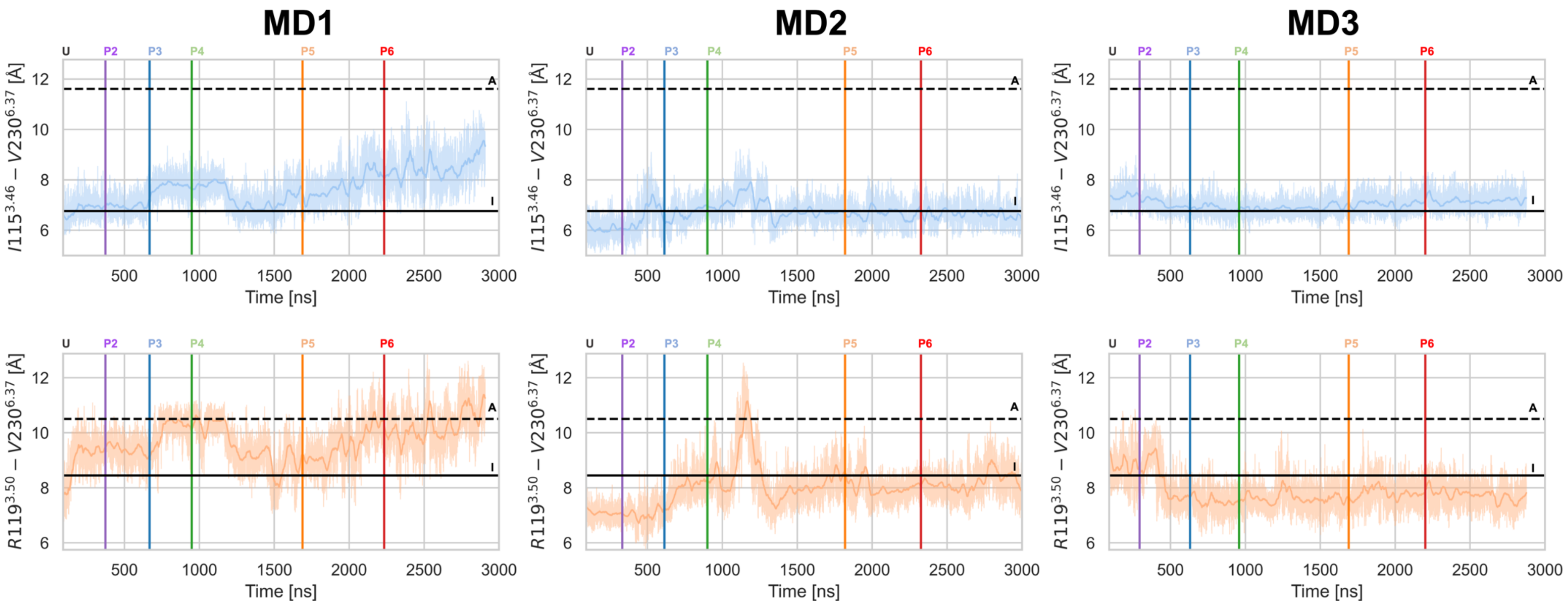


**Figure S8.** Time evolution of distances describing the rewiring of microswitch residue 6.37. The I115$^{3.46}$-V230$^{6.37}$ (blue) and R119$^{3.50}$-V230$^{6.37}$ (orange) distances are shown for the MD1 (left), MD2 (middle) and MD3 (right) pseudo-trajectories. Distances were calculated between the centres of mass of the heavy atoms for each residue pair. Running averages (30-ns window) are shown with a darker hue. Vertical lines separate the individual trajectories with different protonation states (‘U’ to ‘P6’ simulations). The average distance values calculated over all the ‘U’ inactive (solid black line) and the active (dashed black line) state replicas (production only) are shown as horizontal lines.

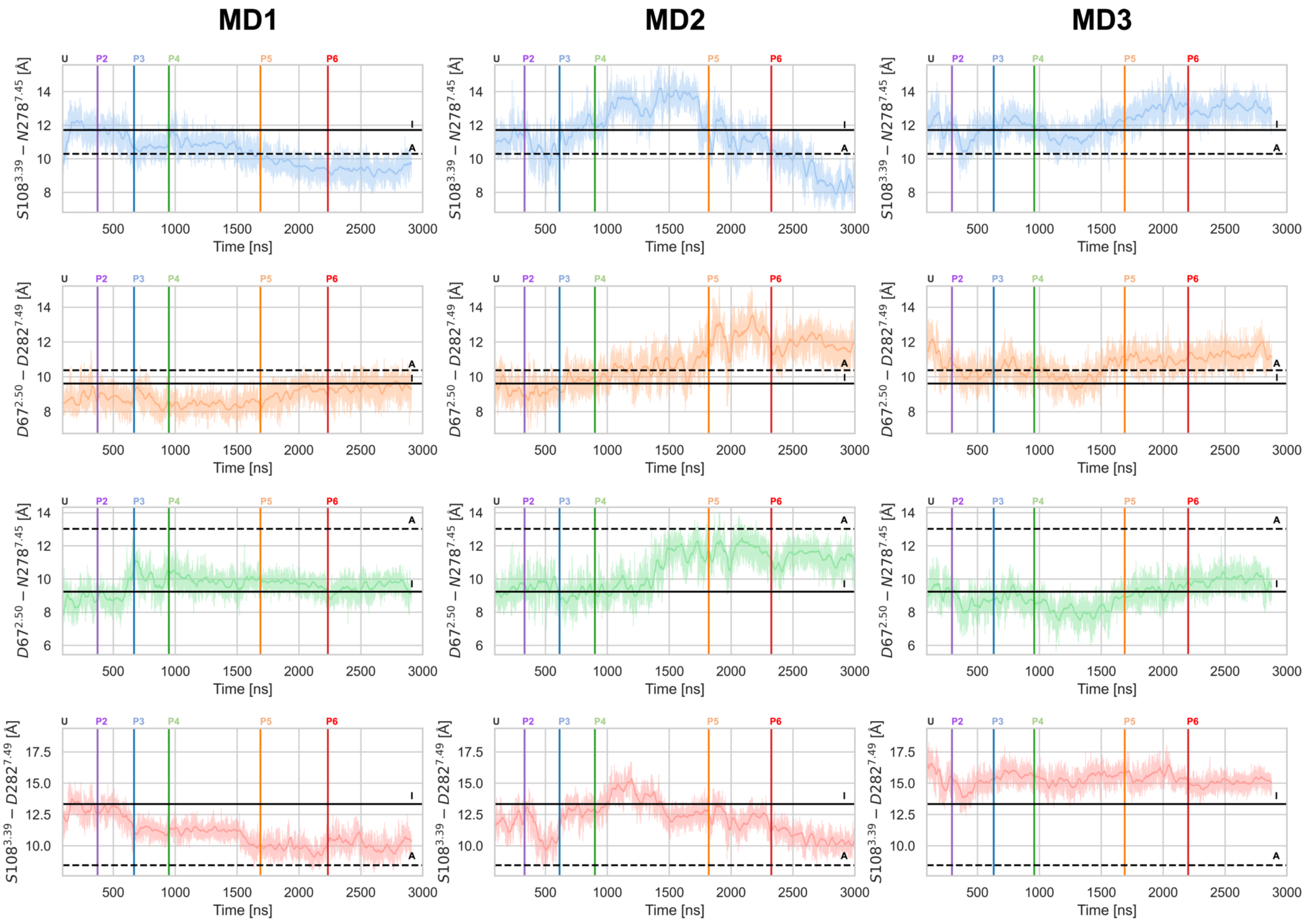

MD1
MD2
MD3
U
P2
P3
P4
P5
P6
I
A
Time [ns]
S108^3.39 – N278^7.45 [Å]
D67^2.50 – D282^7.49 [Å]
D67^2.50 – N278^7.45 [Å]
S108^3.39 – D282^7.49 [Å]

**Figure S9.** Time evolution of distances in the allosteric $Na^+$ binding site. The S108$^{3.39}$-N278$^{7.45}$ (blue), D67$^{2.50}$-D282$^{7.49}$ (orange), D67$^{2.50}$-N278$^{7.45}$ (green), and S108$^{3.39}$-D282$^{7.49}$ (red) distances are shown for the MD1 (left), MD2 (middle) and MD3 (right) pseudo-trajectories. Distances were calculated between the centres of mass of the heavy atoms for each residue pair. Running averages (30-ns window) are shown with a darker hue. Vertical lines separate the individual trajectories with different protonation states ('U' to 'P6' simulations). The average distance values calculated over all the 'U' inactive (solid black line) and the active (dashed black line) state replicas (production only) are shown as horizontal lines.

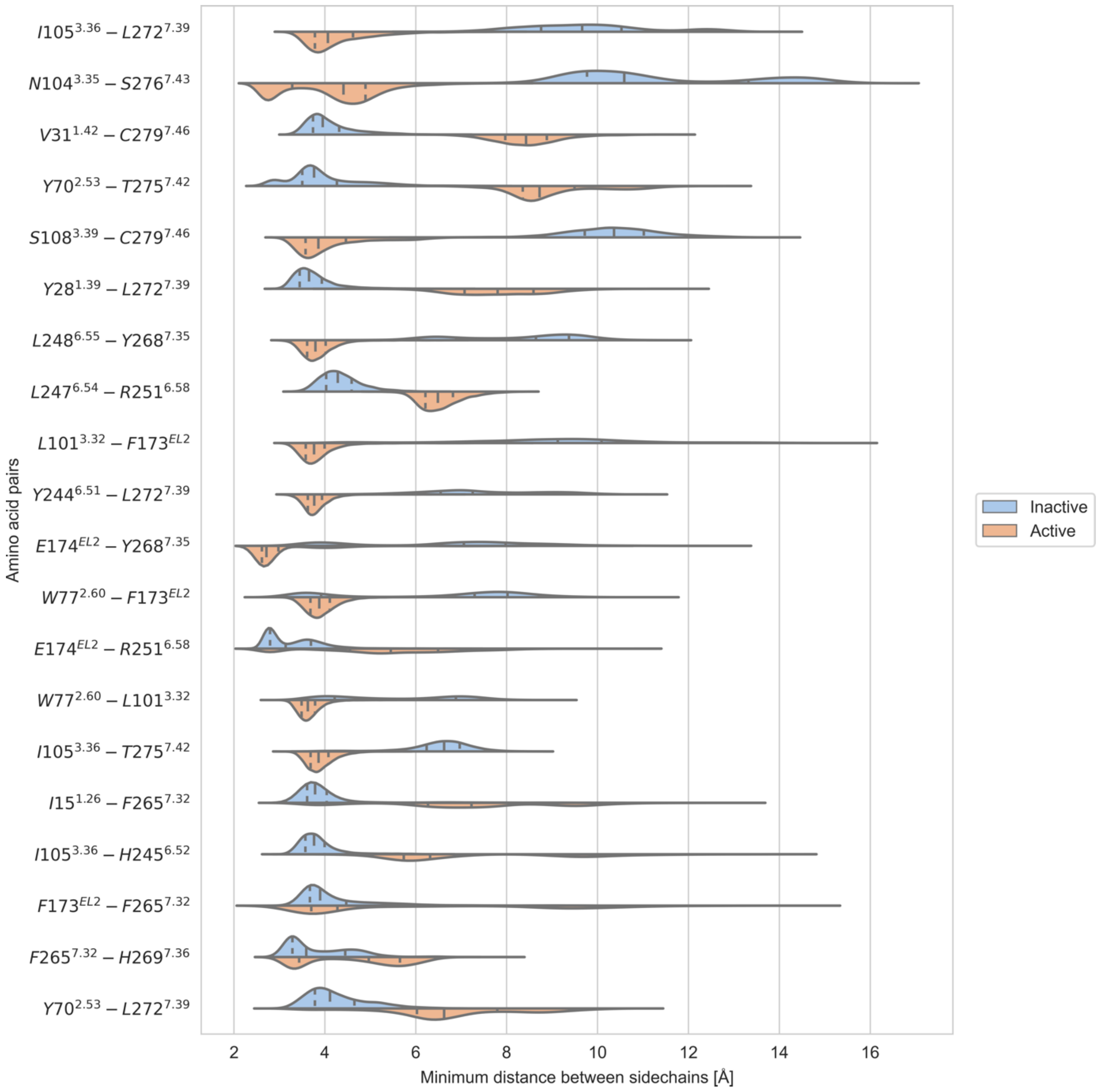


**Figure S10.** Violin plots showing the distribution of distances between the top 20 residue pairs selected through Dynamic Cross-Correlation analysis and sequence data. Minimum distances calculated over all possible pairs of heavy atoms from the side chains of the two residues are shown for the 'U' inactive (blue) and active (orange) replica. The median of each distribution (middle line) and the limits of the two central quartiles (first and third line) are also shown.

8 M. J. Abraham, T. Murtola, R. Schulz, S. Páll, J. C. Smith, B. Hess and E. Lindahl,

GROMACS: High performance molecular simulations through multi-level parallelism from laptops to supercomputers, *SoftwareX*, 2015, **1**–**2**, 19–25.

9 K. Vanommeslaeghe, E. Hatcher, C. Acharya, S. Kundu, S. Zhong, J. Shim, E. Darian, O. Guvench, P. Lopes, I. Vorobyov and A. D. Mackerell, CHARMM general force field: A force field for drug-like molecules compatible with the CHARMM all-atom additive biological force fields, *J. Comput. Chem.*, 2009, NA-NA.

10 X. Yu, X. Huang, T. P. Kenakin, S. T. Slocum, X. Chen, M. L. Martini, J. Liu and J. Jin, Design, Synthesis, and Characterization of Ogerin-Based Positive Allosteric Modulators for G Protein-Coupled Receptor 68 (GPR68), *J. Med. Chem.*, 2019, **62**, 7557–7574.

11 D. W. H. Swenson and S. Roet, Contact Map Explorer, https://contact-map.readthedocs.io/en/latest/index.html.

12 S. Altschul, Gapped BLAST and PSI-BLAST: a new generation of protein database search programs, *Nucleic Acids Res.*, 1997, **25**, 3389–3402.

13 J.-F. Taly, C. Magis, G. Bussotti, J.-M. Chang, P. Di Tommaso, I. Erb, J. Espinosa-Carrasco, C. Kemena and C. Notredame, Using the T-Coffee package to build multiple sequence alignments of protein, RNA, DNA sequences and 3D structures, *Nat. Protoc.*, 2011, **6**, 1669–1682.

14 B. J. Grant, A. P. C. Rodrigues, K. M. ElSawy, J. A. McCammon and L. S. D. Caves, Bio3d: an R package for the comparative analysis of protein structures, *Bioinformatics*, 2006, **22**, 2695–2696.

15 X. P. Huang, T. P. Kenakin, S. Gu, B. K. Shoichet and B. L. Roth, Differential Roles of Extracellular Histidine Residues of GPR68 for Proton-Sensing and Allosteric Modulation by Divalent Metal Ions, *Biochemistry*, 2020, **59**, 3594–3614.

16 J. B. Rowe, N. J. Kapolka, G. J. Taghon, W. M. Morgan and D. G. Isom, The evolution and mechanism of GPCR proton sensing, *J. Biol. Chem.*, 2021, **296**, 100167.

17 J.-P. Liu, T. Nakakura, H. Tomura, M. Tobo, C. Mogi, J.-Q. Wang, X.-D. He, M. Takano, A. Damirin, M. Komachi, K. Sato and F. Okajima, Each one of certain histidine residues in G-protein-coupled receptor GPR4 is critical for extracellular proton-induced stimulation of multiple G-protein-signaling pathways, *Pharmacol. Res.*, 2010, **61**, 499–505.

18 J.-Q. Wang, J. Kon, C. Mogi, M. Tobo, A. Damirin, K. Sato, M. Komachi, E. Malchinkhuu, N. Murata, T. Kimura, A. Kuwabara, K. Wakamatsu, H. Koizumi, T. Uede, G. Tsujimoto, H. Kurose, T. Sato, A. Harada, N. Misawa, H. Tomura and F. Okajima, TDAG8 Is a Proton-sensing and Psychosine-sensitive G-protein-coupled Receptor, *J. Biol. Chem.*, 2004, **279**, 45626–45633.

19 N. Murakami, T. Yokomizo, T. Okuno and T. Shimizu, G2A Is a Proton-sensing G-protein-coupled Receptor Antagonized by Lysophosphatidylcholine, *J. Biol. Chem.*, 2004, **279**, 42484–42491.

20 J. A. Ballesteros and H. Weinstein, Integrated methods for the construction of three-dimensional models and computational probing of structure-function relations in G protein-coupled receptors, *Methods Neurosci.*, 1995, 366–428.
